# Controlling the bioelectrical properties of neurons with ferritin-based Magnetogenetics

**DOI:** 10.1101/2022.12.07.519516

**Authors:** Miriam Hernández-Morales, Koyam Morales-Weil, Sang Min Han, Victor Han, Kelly Pegram, Eric J. Benner, Evan W. Miller, Richard H Kramer, Chunlei Liu

## Abstract

Magnetogenetics promises remote control of neurons, but its validity is questioned due to controversies surrounding the underlying mechanisms and deficits in reproducibility. Recent studies discovered that ferritin, used in Magnetogenetics, transduces radiofrequency (RF) magnetic fields into biochemical signals (reactive oxygen species and oxidized lipids). Magnetic stimulation of ferritin-tethered TRPV channels induces Ca^2+^ responses and modulates behavior but electrophysiological studies indicate that a particular channel, Magneto2.0, is ineffective in affecting the neuronal bioelectrical properties. We investigated this problem using the Magnetogenetic technique FeRIC. We resolved the electromagnetic interference caused by RF in patch-clamp recordings and supported the data with voltage imaging experiments. In neurons expressing TRPV4^FeRIC^, RF depolarizes the membrane potential and increases the spiking frequency. In neurons expressing the chloride-permeable TMEM16A^FeRIC^, RF hyperpolarizes the membrane potential and decreases the spiking frequency. Our study reveals the control of neuronal bioelectrical properties with Magnetogenetics that is non-instantaneous, long-lasting, and moderate, but effective and comparable to that induced by endogenous signaling molecules.

## Introduction

Magnetogenetics aims to manipulate cell activity by using ferritin to transduce magnetic fields into stimuli that activate ion channels. Low to mid-frequency (kHz and MHz) magnetic fields are safe and can freely penetrate through tissues (*1, 2*) allowing wireless, noninvasive, and on-demand control of cell activity. Some magnetogenetic approaches use the transient receptor potential vanilloid 1 and 4 channels (TRPV1 and TRPV4, respectively) coupled with ferritin as magnetic actuators (*3–10*). TRPV1 and TRPV4 are non-selective cation channels activated by heat, pH changes, mechanical stimulus, and a diversity of endogenous lipids (*11–15*). Ferritin, an iron storage protein (*16, 17*), transduces magnetic fields into the stimuli that activate the coupled TRPV channels. However, the mechanism responsible is still disputed. Ferritin has been proposed to transduce static and ultralow frequency magnetic fields (0.08 Hz) into thermal stimulus (*8, 18*) and radiofrequency (RF) magnetic fields (at kHz to MHz frequencies) into biochemical stimuli (*4, 9*). Several studies have reported the magnetic activation of ferritin-coupled TRPV1 and TRPV4 *in vitro* and *in vivo*. Because these channels are permeable to Ca^2+^, their magnetic activation has been examined by monitoring the intracellular Ca^2+^ levels (*3–10*) or by detecting the expression of Ca^2+^-responsive genes (*6, 7*). The magnetic control of neurons has been examined *in vivo* by monitoring the behavioral output from animal models expressing ferritin-coupled TRPV1 or TRPV4 in specific neuronal populations. Magnetogenetics has been used to control escape response and coiling behavior in zebrafish (*10*) and to regulate reward and feeding behavior in mice (*6, 10*). The technique FeRIC has also been used to control the activity of neural crest cells in chicken embryos *in vivo*. The magnetic activation of neural crest cells resulted in neural crest-related craniofacial and heart birth defects. These defects mimicked the effects of endogenous TRPV4 channels activated by fever-like conditions (*3*). Although these studies indicate that magnetic fields activate ferritin-coupled ion channels, their effects on the membrane electrical properties remains unknown, further complicating the development of Magnetogenetics.

One study reported that stimulating neurons expressing Magneto2.0, a ferritin-fused TRPV4, in the entorhinal cortex (*ex vivo*) and the striatum (*in vivo*) with static magnetic fields triggers action potential firing (*10*). However, a series of independent studies using *in vitro* and *in vivo* patch-clamp and extracellular electrophysiological (Ephys) recordings reported that magnetic stimulation failed to trigger action potential firing and did not change the electrical properties in multiple types of neurons expressing Magneto2.0 (*19–21*). The negative results from experiments using Magneto2.0 add uncertainty to the field of Magnetogenetics. Moreover, there is no Ephys characterization of other ferritin-coupled ion channels, such as the αGFP-TRPV1/GFP-ferritin (*6*) or the FeRIC channels (*3–5*), which have been reported to be activated with RF. This is, in part, due to the technical challenge caused by the electromagnetic interference between the RF setup and the Ephys system. Here, we solved that problem and combined Ephys recordings with voltage and calcium imaging to characterize the RF control of the membrane potential in Neuro2a (N2a) cells and cultured hippocampal neurons. We first characterized the inward currents and the membrane depolarization induced by RF stimulation of cells expressing TRPV4^FeRIC^. We then characterized the chloride influx and the subsequent membrane hyperpolarization produced by RF stimulation of the new FeRIC channel TMEM16A^FeRIC^, which is permeable to chloride ions (Cl^-^). Using FeRIC, we show that RF activates the ferritin-coupled TRPV4 and TMEM16A channels, producing opposite effects on the membrane potential and cell activity, thus supporting Magnetogenetics as a valid and viable technique for modulating membrane bioelectrical properties.

## Results

### RF decreases the membrane resistance and produces inward currents in cells expressing TRPV4^FeRIC^

To solve the electromagnetic interference problem between the Ephys system and the magnetic field stimulus, we used of a relatively high RF magnetic field frequency stimulus. Because of the nearly five orders of magnitude difference in frequencies between our RF system (at 180 MHz) and the Ephys setup (lowpass-filtered at 3 kHz), low electromagnetic interference was expected. However, when implemented, RF consistently interfered with the Ephys setup (**Fig. 1**). The RF stimulus couples to the Ephys setup, and through Faraday’s law, the RF magnetic fields produce electric fields on the wires, which ultimately produce RF voltages on the patch-clamp headstage. The amplitudes of the electric fields and thus voltages are linearly proportional to the rate of change of magnetic flux through the loop(s) formed by the various Ephys circuit elements (wires, electrodes, cell bath, etc). The RF voltages induced do not show up directly on the patch-clamp recordings because they are orders of magnitude higher in frequency. Instead, they appear in the form of baseline shifts due to nonlinear effects (**Fig. 1**). When implementing the Ephys setup with our previously reported RF system (5 cm RF coil diameter) (*5*), RF at 180 MHz and 6 μT produced a baseline shift of the measured current from a pipette in-the-bath of more than 4400 pA (**Fig. 1B**). This baseline current shift is well above the average amplitude of a neuronal action current, hindering the feasibility of Ephys recordings of neurons under RF stimulation. To reduce the magnetic flux, a custom-built RF emitting coil (2.8 cm in diameter) was attached to the bottom of the recording chamber. The small RF coil allows focused delivery of the RF stimulus near the region of the recording chamber containing the cells (**Fig. 1C – E**) and decreases the electrical interference transmitted to the patch-clamp electrode by up to 50% compared with the 5 cm diameter coil we used in previous studies (**Fig. 1B**) (*5*). The 180 MHz voltages induced on the head stage input impedance due to the electromagnetic fields were simulated to be less for the smaller coil for a variety of electrode wire geometries (**Fig. 1C, D**). To further decrease the RF interference, two ferrite beads designed to block frequencies near 180 MHz were placed over the cables connected to the RF coil (**Fig. 1A**). These ferrite beads prevented unwanted RF radiation from the cables and drastically reduced the minimal baseline current shift under RF stimulation (**Fig. 1B**). Next, using the 2.8 cm diameter RF coil and the ferrite beads, we measured the basal shift in the current and the voltage of a pipette in-the-bath produced by RF at 180 MHz ranging from 0.2 to 10 μT in voltage-clamp (holding voltage = 0 mV) and current-clamp (holding current = 0 pA) configurations (**Fig. 1F, G**). The baseline shifts in both configurations as a function of RF magnetic field amplitude are fit with a quadratic term. Because of the dependence on magnetic flux, loop area, and other circuit properties, this is a geometry-dependent problem. To isolate the problem from the geometry, verify the effect, and to measure its characteristics in a repeatable manner, the baseline shift was also measured as a function of the RF voltage produced and directly set by an arbitrary waveform generator directly connected or capacitively coupled to the headstage. We measured the baseline shift in voltage-clamp and current-clamp configurations at the two RF frequencies, 180 MHz and 345 kHz (**Fig. 1H, I**), reported to activate TRPV1 and TRPV4 channels coupled with ferritin (*3–5, 22, 23*). For low RF voltages, the baseline shifts (pA) in the voltage-clamp configuration as a function of RF voltage are fit with a quadratic term (Fig. 1D, inset). As the RF voltage and corresponding baseline shifts increase, higher-order nonlinearities dominate over the initial second-order nonlinearity, making the baseline shifts increase even faster. Remarkably, 345 kHz RF saturates the patch-clamp amplifier at a voltage over an order of magnitude less than that of 180 MHz RF; for small RF voltages, the coefficient of the quadratic fit is orders of magnitude larger for 345 kHz than 180 MHz (**Fig. 1H**). The baseline shifts (mV) in the current-clamp configuration as a function of RF voltage are fit with quadratic terms with the 345 kHz data having a higher coefficient relative to 180 MHz (**Fig. 1I**). These results emphasize the need to keep RF voltages induced on the headstage low and that RF at 345 kHz produces larger baseline shifts compared with RF at 180 MHz. Therefore, as we did in our previous studies, we conducted all the experiments using RF at 180 MHz and 1.6 μT. With the RF voltages induced on the headstage being proportional to the RF magnetic field strength, a larger RF magnetic field strength was not used because it produces a non-negligible shift in the basal current level.

**Fig. 1.**
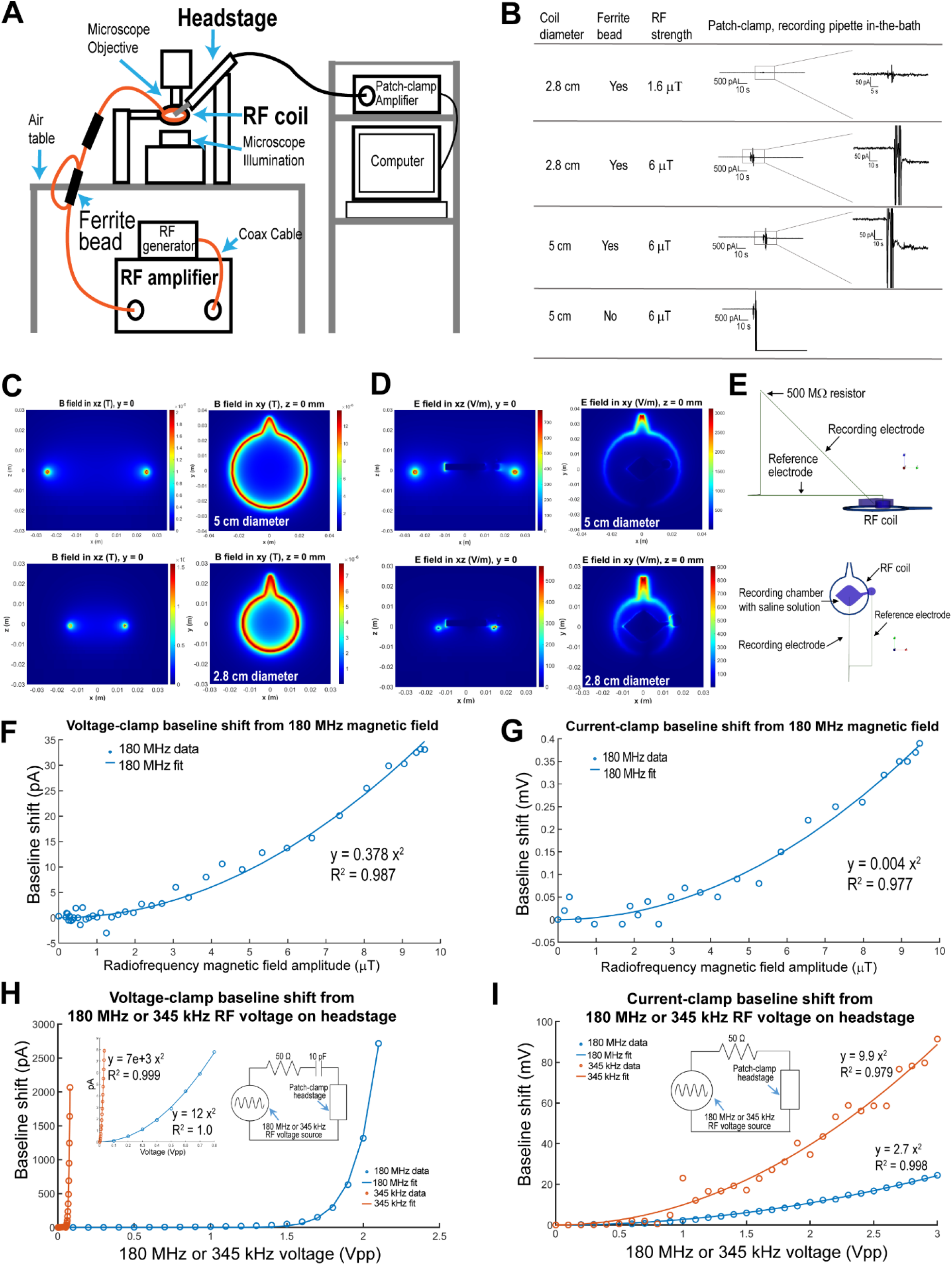
Electromagnetic interference between the Ephys setup and the RF system emitting at 180 MHz and 345 kHz. (**A**) Illustration of the Ephys setup and RF system used for patch-clamp recordings of cells stimulated with RF. (**B**) Shifts of the baseline current recorded from a pipette in-the-bath (6.9 MΩ) in the voltage-clamp configuration (0 mV). For each RF condition, the RF emitting coil diameter, the use of ferrite beads, and the RF amplitude are indicated. (**C, D**) Finite-difference time-domain electromagnetic simulation results are shown for an RF coil of 5 cm or 2.8 cm diameter. The magnitudes of 180 MHz (**C**) magnetic and (**D**) electric fields are shown with all fields scaled for a magnetic field with an amplitude of 1.6 µT at the center. (**E**) Metal for the RF coils and patch-clamp electrodes are modeled as well as a two-compartment conductive saline solution bath and a patch-clamp head stage input impedance (500 MΩ). The amplitudes of the electric fields at the center were 13.1 V/m and 5 V/m for the large RF coil and the small RF coil, respectively. (**F, G**) The shift of the baseline recorded from a pippete in-the-bath (8 MΩ) is shown for the (**F**) voltage-clamp or (**G**) current-clamp configuration as a function of the 180 MHz RF amplitude in μT. The data fit (solid lines) and the equations are indicated. (**H, I**) The baseline current shift is shown for the (**H**) voltage-clamp (inset, zoom in) or (**I**) current-clamp configuration as a function of RF voltage on the voltage source from the RF generator at 180 MHz (blue) or 345 kHz (orange). The data fit (solid lines) and the equations are indicated. Insets: Illustrations of the setups used to measure the baseline shifts produced by an RF source at 180 MHz or 345 kHz.

To prevent bias and error, even under minimal RF interference, we measured the membranes’ electrical properties before RF (Baseline), during RF stimulation (180 MHz and 1.6 µT), and within the 2 minutes after RF stimulation (Post-RF). Because the mechanism underlying the RF-induced activation of FeRIC channels is biochemical and sustains after turning off the RF, the RF-induced effects can be observed for a few minutes after RF is turned off (*5*). To corroborate that RF activates TRPV4^FeRIC^ and affects the membrane’s properties, we monitored the membrane resistance (Rm), by conducting patch-clamp experiments in the whole-cell, voltage-clamp configuration (Vh = -60 mV) (**Fig. 2**). A decrease in Rm indicates an increase in the number of open ion channels at the cell membrane. The Rm was assessed by applying a 5 mV square voltage pulse every 30 seconds. This strategy offers the advantage of simultaneously monitoring the Rm and the access resistance (Ra) to rule out the possibility of observing changes in Rm due to changes in the patch-clamp seal quality. Experiments included in the analysis are those where Ra did not change more than 20% with respect to the baseline during the span of the experiment. In N2a cells expressing TRPV4^FeRIC^, RF stimulation for 5 minutes decreased the Rm to 61.5 ± 8.5 % (n = 13 cells, p < 0.05) relative to the baseline (**Fig. 2A, G**). This effect was not observed in the presence of the TRPV4 antagonist GSK 2193874 (GSK219 at 1 µM; 88.5 ± 8.1 %; n = 8 cells, p = 0.38; **Fig. 2B, G**), suggesting that the decrease in the Rm is due to the activation of TRPV4^FeRIC^. To test if RF acts on native ion channels, we applied RF to mock-transfected N2a cells. In those cells, RF did not change the Rm (95.5 ± 10.3 %; n = 6 cells, p = 0.71; **Fig. 2C, G**). Similarly, in N2a cells expressing the non-conductive TRPV4^FeRIC^ mutant (M680D/ΔK675), RF stimulation did not affect the Rm (111.5 ± 4.2 %; n = 8 cells, p = 0.68; **Fig. 2D, G**). Finally, the RF effects on the membrane’s electric properties were replicated with the TRPV4 agonist GSK1016790A (GSK101), which decreased the Rm to 60.7 ± 12.3 % (n = 6, p < 0.05) relative to the baseline. This effect was inhibited with GSK219 (113.0 ± 11.6%; n = 10 cells; **Fig. 2E** – **G**).

**Fig. 2.**
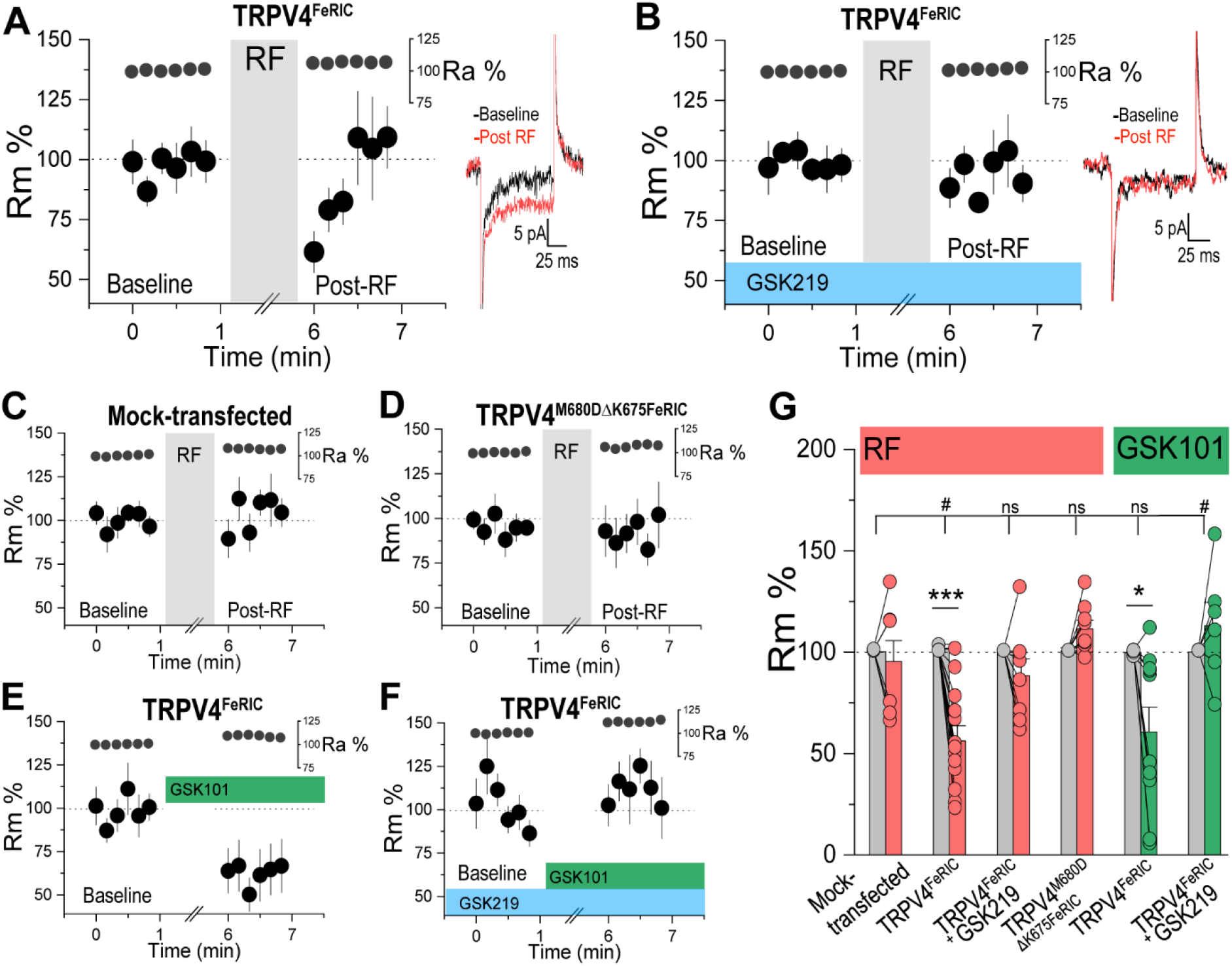
RF decreases the membrane resistance in Neuro2a cells expressing TRPV4^FeRIC^. (**A** – **F**) Time course of the average changes (±SEM) in the membrane resistance (Rm) and access resistance (Ra) from N2a cells recorded in the voltage-clamp mode before and following stimulation with RF or GSK101. Rm and Ra (average ± SEM) from N2a cells expressing TRPV4^FeRIC^ before and following RF stimulation (5 min, gray box) in (**A**) the absence or (**B**) the presence of the TRPV4 antagonist GSK219. Insets: representative traces of whole-cell currents in the baseline (black) and in the post-RF period (red). Rm and Ra (average ± SEM) from (**C**) mock-transfected N2a cells or (**D**) N2a cells expressing the nonconductive TRPV4^FeRIC^ mutant (M680D/ΔK675) before and following RF stimulation. Average changes (±SEM) in Rm and Ra from N2a cells expressing TRPV4^FeRIC^ before and following the application of the TRPV4 agonist GSK101 in (**E**) the absence or (**F**) the presence of GSK219. In experiments with GSK219, cells were treated at least 10 min before applying RF or GSK101. (**G**) Average of Rm (±SEM) in N2a cells from the different experimental groups. Significance was determined using a two-tailed Student’s t-test (baseline vs. RF or GSK101 comparisons) and a two-way ANOVA followed by Holm-Sidak multiple comparisons test. Where applicable, p < 0.05 (∗) or p < 0.0001 (∗∗∗) was considered a statistically significant difference.

Next, to assess the effects of RF on membrane currents, we patch-clamped (Vh = -60 mV) N2a cells expressing TRPV4^FeRIC^ and measured the mean amplitude in pA, the total time of the activated inward currents in seconds, and estimated the charges carried (see Methods). In baseline conditions, N2a cells expressing TRPV4^FeRIC^ showed infrequent inward currents with an amplitude of 3.6 ± 0.25 pA, total time duration of 8.5 ± 2.3s, and a total carried charge of 33.1 ± 9.1 pA•s (n = 6 cells) (**Fig. 3A, B**). Compared to the baseline, RF (180 MHz and 1.6 μT for 5 min) increased these inward currents parameters (3.7 ± 0.25 pA, 9.4 ± 2.1 s, 36.6 ± 8.3 pA•s; n = 6 cells) and reached a statistically significant difference in the post-RF period (Post-RF: 4.1 ± 0.27 pA, 14.4 ± 3.4 s, 58.8 ± 12 pA•s; p < 0.05; n = 6) (**Fig. 3A, B**). The RF effects on the membrane currents were blocked by GSK219 (n = 7) (**Fig. 3C, D**). The RF-induced currents mediated by TRPV4^FeRIC^ were transient, with varying durations, and returned to the basal level approximately 5 minutes after RF stimulation was turned off. These observations suggest that RF does not activate all TRPV4^FeRIC^ channels simultaneously and that the RF-produced stimuli are not cleared immediately after RF is switched off. Remarkably, the inward currents mediated by TRPV4^FeRIC^ are similar to those observed when the native TRPV4 is activated by endogenous lipids (*11*). To estimate the unitary conductance of TRPV4^FeRIC^, we performed a nonstationary noise analysis (see Methods) of the whole-cell inward currents recorded in N2a cells expressing TRPV4^FeRIC^. In this analysis, the relationship between the mean current amplitude (*I*) and the variance of the currents (σ^2^) is used to estimate the ion channels’ unitary current (*i*) and unitary conductance (γ) (**Fig. 3E**) (*24–28*). In our experimental conditions, the estimated *i* for TRPV4^FeRIC^ was -2.2 ± 0.22 pA in the baseline and - 2.3 ± 0.26 pA in the post-RF period (**Fig. 3F**). Using these values, the estimated γ of TRPV4^FeRIC^ was 34 ± 3.48 pS in the baseline and 35 ± 4.1 pS in the post-RF period (**Fig. 3G**). The estimated TRPV4^FeRIC^ conductance (γ) is similar to that reported for the wild-type TRPV4 which range from 30 – 60 pS (*29, 30*). We did not observe differences when comparing the *i* or γ of TRPV4^FeRIC^ before or after RF stimulation (p = 0.79, n = 6 cells) (**Fig. 3F, G**), suggesting that the RF-induced inward currents are due to an increase in the open channel probability of TRPV4^FeRIC^. To corroborate that RF does not affect the basal electrophysiological properties of N2a cells, we measured the relationship between voltage and currents (V-I) in baseline conditions (preceding RF stimulation) and 5 minutes after RF stimulation. In mock-transfected N2a cells (n = 6 cells, **Fig. S1A**) and N2a cells expressing TRPV4^FeRIC^ (n = 10 cells, **Fig. S1B**) the V-I relations before and after RF stimulation were not significantly different. To examine if RF generates concurrent inward currents and cytosolic Ca^2+^ increases, we conducted simultaneous patch-clamp in the whole-cell, voltage-clamp configuration and Ca^2+^ imaging experiments. In N2a cells expressing TRPV4^FeRIC^ and gCaMP6, RF stimulation produced inward currents and cytosolic Ca^2+^ increases (**Fig. S2**). The RF-induced Ca^2+^ increases in N2a cells were similar to those observed in our previous studies (*3*– *5*). In general, the inward currents preceded the cytosolic Ca^2+^ increases, but the Ca^2+^ increases lasted longer (**Fig. S2B, C**). The Ephys and Ca^2+^ imaging results indicate that RF activates TRPV4^FeRIC^ generating inward currents, decreasing the Rm, and increasing the cytosolic Ca^2+^ levels.

**Fig. 3.**
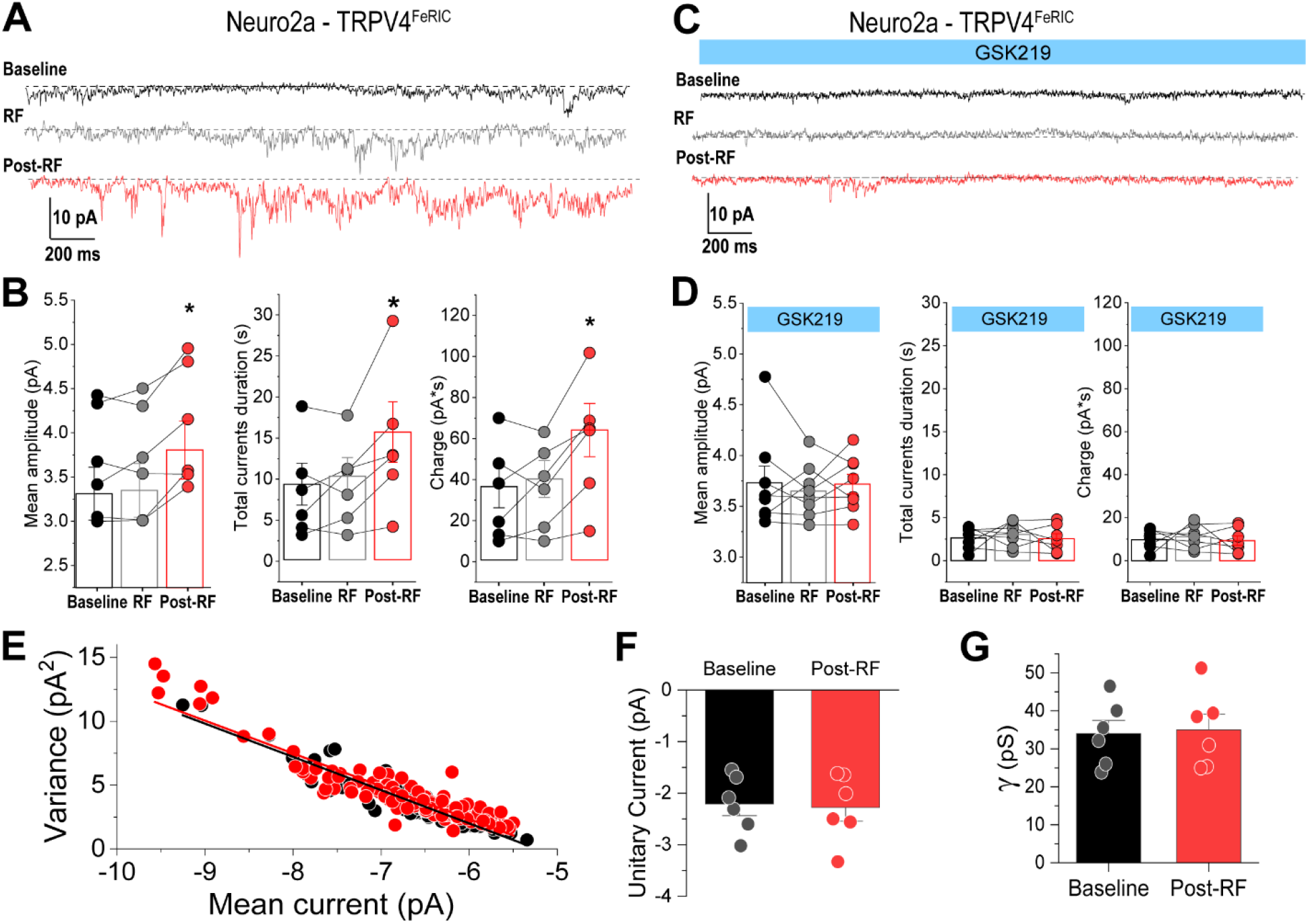
RF evokes inward currents in Neuro2a cells expressing TRPV4^FeRIC^. (**A**) Representative whole-cell currents on voltage-clamp before RF (baseline, black), during RF (180 MHz and 1.6 μT, gray), and within the 2 minutes after RF stimulation (post-RF, red) in an N2a cell expressing TRPV4^FeRIC^. (**B**) Mean current amplitude, total currents duration, and the estimated carried charge in the baseline, during RF, and in the post-RF period. (**C**) In a different set of experiments, cells were treated with GSK219. Membrane currents were recorded in the baseline, during RF, and in the post-RF period. (**D**) Mean current amplitude, current duration, and carried charge in the baseline, during RF, and in the post-RF period in cells treated with GSK219. (**E**) Representative current-variance relationships during baseline (black) and in the post-RF (red) period. Data were fit using the equation for the non-stationary noise analysis (see the Methods section). (**F**) Average unitary current (*i*) and (**G**) unitary conductance (γ) in the baseline and in the post-RF period. Significance was determined using a two-tailed Student’s t-test. Where applicable, p < 0.05 (∗) was considered a statistically significant difference.

### RF depolarizes hippocampal neurons expressing TRPV4^FeRIC^

To examine the control of neuronal membrane potential with FeRIC, we conducted Ephys experiments in cultured hippocampal neurons expressing TRPV4^FeRIC^, identified as those expressing mCherry (mCherry+). First, we verified that TRPV4^FeRIC^ expression does not affect the basal passive and active electrical properties of neurons. We applied square voltage steps to obtain the I-V relation and measured the number and amplitude of the action potentials as well as the instantaneous frequency (**Fig. S1C–-E**). All these parameters were similar in mock-transfected neurons (n = 8 cells) and neurons expressing TRPV4^FeRIC^ (n = 11; **Fig. S1D, E**). Next, we monitored the membrane potential before RF (Baseline), during RF (5 min, 180 MHz, 1.6 μT), and within 2 minutes after RF stimulation (Post-RF). In mock-transfected neurons, RF did not affect the membrane potential (Basal: -59.9 ± 1.8 mV, Post-RF: -60.1 ± 2.1 mV; p = 0.78; n = 5 cells**; Fig. 4A, B**), but in neurons expressing TRPV4^FeRIC^, RF gradually depolarized the membrane potential (Basal: -58.8 ± 1.8 mV, Post-RF: -56.6 ± 1.6 mV; p < 0.05; n = 7 cells; **Fig. 4C, D**). The antagonist GSK219 inhibited the RF-induced membrane depolarization (Baseline: -59.14 ± 1.68 mV, RF: - 58.65 ±0.86 mV; p = 0.66; n = 8 cells; **Fig. 4E, F**). Consistently, in neurons expressing TRPV4^FeRIC^, RF decreased the Rm with respect to the basal level (80 ± 10 %; n =7 cells; p < 0.05; **Fig. 4D inset**), but did not change the Rm in neurons treated with GSK219 (103 ± 9%; n = 8; **Fig. 4F inset**) or in mock-transfected neurons (101 ± 6%; n = 5; **Fig. 4B inset**). Our results are in contrast to previous studies that found static magnetic stimulation evoking immediate action potential firing in neurons expressing ferritin-tagged TRPV1 (*6*) and ferritin-tagged TRPV4 (*10*). We did not observe this instantaneous effect using the FeRIC system. To examine the consequences of the RF-induced membrane depolarization in a larger number of neurons, we conducted voltage imaging experiments with the voltage-sensitive fluorophore BeRST 1 (*31, 32*). We resolved the neuronal electrical activity on a millisecond time scale. Hippocampal neurons expressing either TRPV4^FeRIC^ or TRPV4^WT^ were labeled with BeRST 1 (500 nM) and imaged at 46.5 Hz (**Fig. 4G, H**). The slow imaging rate did not yield the temporal resolution needed to optically resolve individual action potentials. Therefore, we developed a custom algorithm tailored to inferring the fast membrane depolarization events or spikes from the time series data of BeRST 1 fluorescence changes (see Methods). The algorithm takes as input the BeRST 1 fluorescence change time series data and outputs binary spikes. Although we were not able to definitely distinguish between action potentials, calcium spikes, or postsynaptic potentials, examining the BeRST 1 ΔF/F0 changes under this setting led to consistent observations of the neuronal spiking activity, and we were able to observe several different spiking activity patterns (**Fig. S3**). The reported BeRST 1 data are the average spiking frequency (Hz ± SEM), and the data include both spiking and non-spiking neurons. BeRST 1-stained neurons were imaged with no added antagonists of neurotransmitter receptors or blockers of ion channels. Neurons were imaged in the absence of RF stimulation (No RF) and during RF stimulation (from 30 to 120 s; **Fig. 4H**). In TRPV4^FeRIC^-expressing neurons (mCherry+), RF stimulation significantly increased the spike frequency compared to that of mock-transfected neurons or non-stimulated TRPV4^FeRIC^-expressing neurons (Mock-transfected + RF: 0.006 ± 0.002 Hz; TRPV4^FeRIC^ + NO RF: 0.13 ± 0.06 Hz; TRPV4^FeRIC^ + RF: 0.34 ± 0.09 Hz; n = 81 – 181 cells; p < 0.05). This effect was abolished by the antagonist GSK219 (0.03 ± 0.02 Hz; n = 22 cells; p = 0.08) (**Fig. 4I**). As expected, RF did not increase the spiking frequency in neurons expressing the wild-type TRPV4 (0.02 ± 0.01 Hz; n = 15 cells) (**Fig. 4I**). RF also increased the percent of TRPV4^FeRIC^-expressing neurons that displayed spiking activity (**Fig. 4K**). Interestingly, during RF stimulation, neighboring neurons that did not express TRPV4^FeRIC^ (mCherry-) also increased the spiking frequency (No RF: 0.08 ± 0.03 Hz, n = 205 cells; RF: 0.21 ± 0.05 Hz; n = 229 cells; p < 0.0001). These results indicate that RF depolarizes the membrane potential and increases the neuronal spiking activity in both neurons expressing TRPV4^FeRIC^ and the neighboring neurons not expressing TRPV4^FeRIC^.

**Fig. 4.**
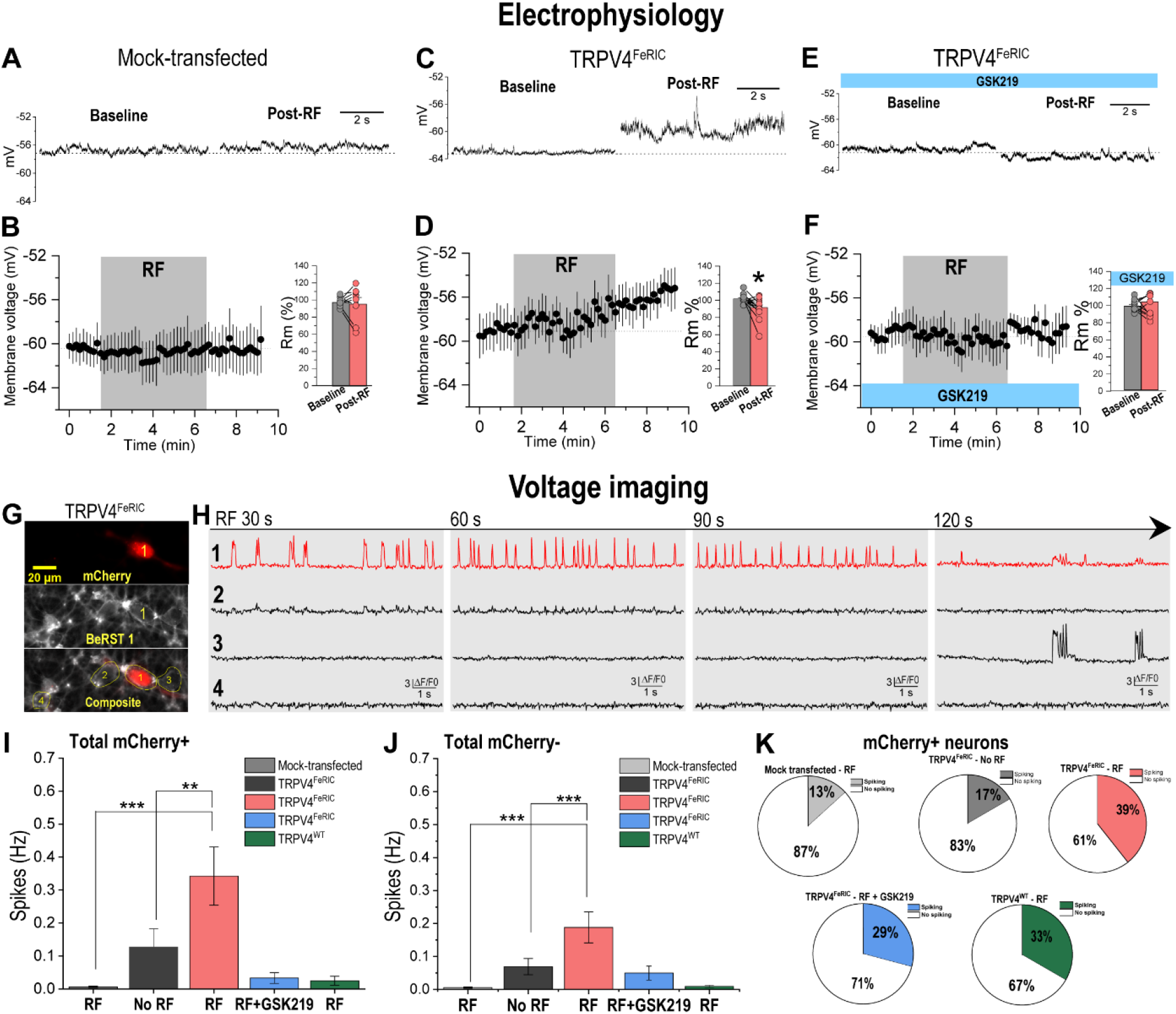
RF depolarizes the membrane potential and increases the spiking frequency in hippocampal neurons expressing TRPV4^FeRIC^. (**A** - **F**) Membrane voltages recorded from neurons in the current-clamp mode. Representative traces of the membrane voltage before (baseline) and within the 2 minutes after RF stimulation (post-RF) in (**A**) mock-transfected neurons or neurons expressing TRPV4^FeRIC^ in (**C**) the absence or (**E**) the presence of GSK219. Time course of the average membrane potential (±SEM) before RF, during RF, and in the post-RF period in (**B**) mock-transfected neurons or neurons expressing TRPV4^FeRIC^ (**D**) in the absence or (**F**) in the presence of GSK219. Insets: average Rm (±SEM) in the baseline and the post-RF period. (**G**) Epifluorescence images of neurons expressing TRPV4^FeRIC^ (mCherry^+^) stained with BeRST 1. (**H**) Changes in BeRST 1 fluorescence in the mCherry+ neuron (1) and the neighboring mCherry-neurons (2 – 4) from (G). After 30 s of RF stimulation (gray box), BeRST 1 fluorescence was acquired for consecutive periods of 10 s. Averages (±SEM) of spikes (Hz) for (**I**) total mCherry^+^ neurons (spiking and non-spiking) and for (**J**) total mCherry-neurons (spiking and non-spiking). Experimental groups are independent samples. (**K**) Pie charts of the fraction of mCherry+ neurons that were spiking (at least one spike in the entire experiment) or not spiking for the different experimental groups. For patch-clamp results, significance was determined using a two-tailed Student’s t-test. For BeRST 1 results, significance was determined using a Kruskal-Wallis ANOVA followed by Dunn’s multiple comparisons test. Where applicable, p < 0.05 (∗), p < 0.001 (∗∗), or p < 0.0001 (∗∗∗) was considered a statistically significant difference.

### RF activates the chloride channel TMEM16A^FeRIC^

In the central nervous system, inhibitory transmission is mediated primarily by chloride (Cl^-^) currents through diverse ligand-gated ionotropic receptors such as GABA and glycine receptors. Here, we designed a new FeRIC channel permeable to Cl^-^ by fusing the anoctamin 1 (ANO1), also known as the transmembrane member 16A (TMEM16A), with the domain 5 of kininogen at the amino-terminal. TMEM16A is activated with cytosolic Ca^2+^ increases, cell swelling and stretch, noxious temperatures (∽ 44°C), acidic pH, ROS, and diverse lipids and lipid peroxidation products(*33–35*). Because TMEM16A is sensitive to ROS and lipids, we expected RF to activate this channel. To examine the RF-induced activation of TMEM16A^FeRIC^, we monitored the Rm in cells expressing FeRIC and wild-type TMEM16A, before RF (baseline) and in the 2-minute time window after RF stimulation. In N2a cells expressing TMEM16A^FeRIC^, RF decreased the Rm to 62.8 ± 10.4 % (n = 6 cells; p < 0.05) relative to the baseline (**Fig. 5A, B**). This effect was not observed in cells expressing TMEM16A^WT^ (104 ± 10 %; n = 5 cells; p = 0.86; **Fig. 5C, D**), suggesting that the change in the Rm is due to the activation of TMEM16A^FeRIC^. To corroborate that RF activates TMEM16A^FeRIC^, we used the genetically encoded halide indicator YFP-H148Q (YFP). The fluorescence of YFP is quenched by halide ions, but YFP exhibits the maximum sensitivity to iodide (*36*). Because TMEM16A is permeable to both chloride and iodide, we used iodide to monitor its activation. Responses were quantified as the ΔF/F0 of YFP fluorescence where F0 was measured during the first 60 seconds prior to iodide application. In TMEM16A^FeRIC^-expressing N2a cells, RF stimulation for 5 minutes significantly decreased the YFP fluorescence (−12.7 ± 1 ΔF/F0; n=3, 198 cells) compared to that in non-stimulated cells (0.003 ± 0.6 ΔF/F0; n=7, 411 cells; p < 0.0001). The TMEM16A antagonist T16Ainh-A01 (T16Ainh, 2 µM) significantly inhibited the RF-induced decrease in YFP fluorescence (−5.2 ± 0.8 ΔF/F0; n=3, 176 cells; p < 0.0001) (**Fig. 5E, F**). In contrast, RF stimulation of TMEM16A^WT^-expressing cells did not change the YFP fluorescence (−0.5 ± 1 ΔF/F0; n=3, 171 cells) when compared to non-stimulated cells (0.2 ± 1 ΔF/F0, n=3, 172 cells; p = 0.84) (**Fig. 5G, H**). Because TMEM16A is activated with the rise in cytosolic Ca^2+^, the functional expression of TMEM16A^FeRIC^ and TMEM16A^WT^ was corroborated with ionomycin, a Ca^2+^ ionophore. Ionomycin decreased the YFP fluorescence in cells expressing TMEM16A^FeRIC^ (−23.9 ± 1 ΔF/F0, n=6, 377 cells) and TMEM16A^WT^ (−11.9 ± 1 ΔF/F0, n=3, 336 cells). This effect was inhibited with T16Ainh (FeRIC: -10.1 ± 0.68 ΔF/F0, n=4, 317 cells; wild-type: -4.5 ± 0.9 ΔF/F0; n=5, 371 cells; p < 0.001) (**Fig. 5E, F, H**). These results show that coupling endogenous ferritin with TMEM16A enables RF to control its activation.

**Fig. 5.**
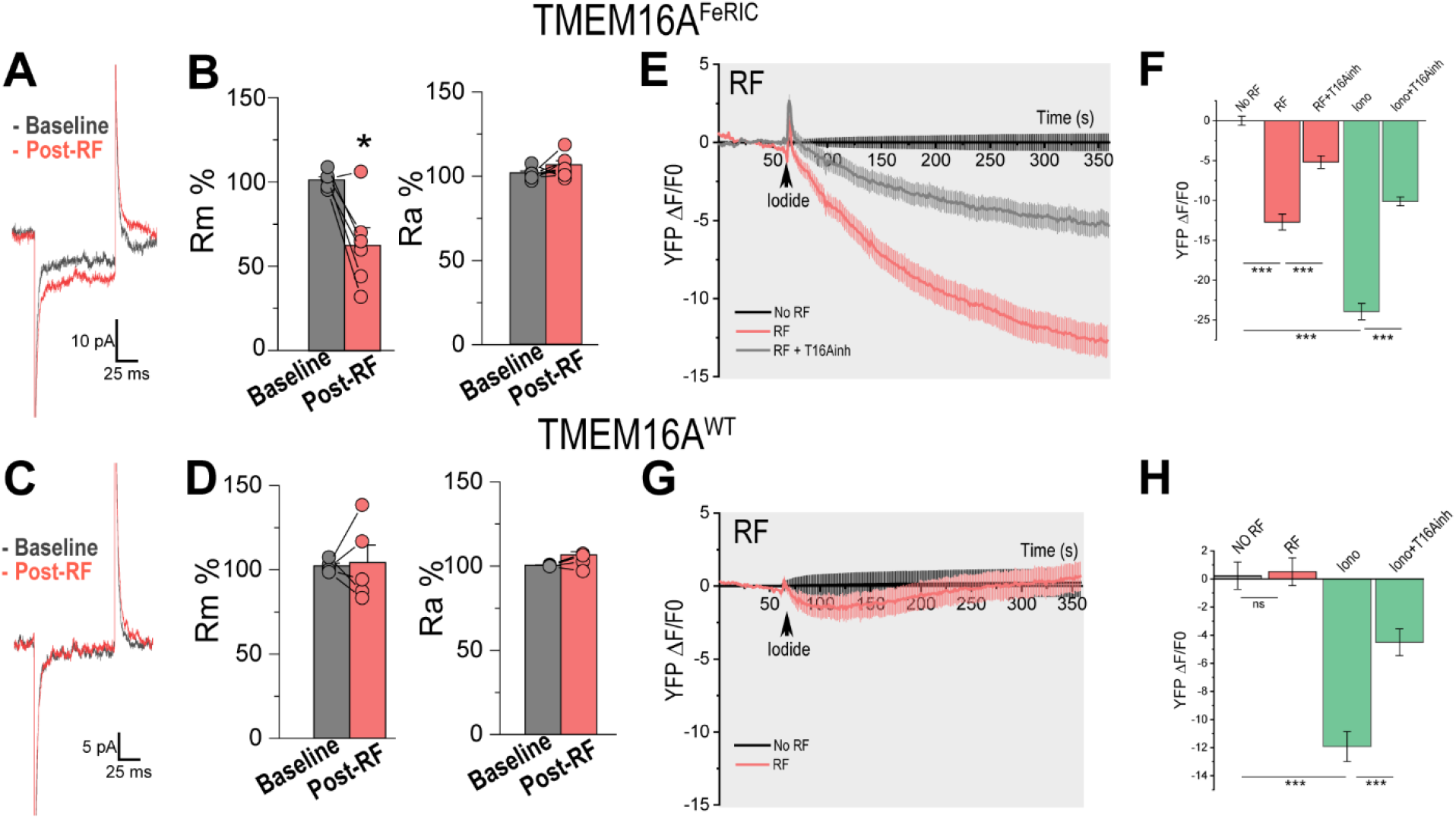
RF decreases the membrane resistance and increases the anion influx in Neuro2a cells expressing TMEM16A^FeRIC^. (**A** - **D**) Electrical properties of N2a cells expressing TMEM16A^FeRIC^ and TMEM16A^WT^ were recorded in the voltage-clamp mode. Representative traces of whole-cell currents from N2a cells expressing (**A**) TMEM16A^FeRIC^ or (**C**) TMEM16A^WT^ in the baseline (gray) and within the 2 minutes after RF stimulation (post-RF, red). Average changes (±SEM) in Rm and Ra in the baseline and the post-RF period from N2a cells expressing (**B**) TMEM16A^FeRIC^ or (**D**) TMEM16A^WT^. Average changes (±SEM) in YFP-H148Q (YFP) fluorescence in N2a cells expressing (**E**) TMEM16A^FeRIC^ or (**G**) TMEM16A^WT^ following the addition of imaging solution containing 70 mM iodide (black arrow) in the absence (black) or following RF stimulation (red). In a different set of experiments, cells were treated with the TMEM16A antagonist T16Ainh-A01 (T16Ainh, gray). Averages (±SEM) of the change in YFP fluorescence after 5 min of iodide application in N2a cells expressing (**F**) TMEM16A^FeRIC^ or (**H**) TMEM16A^WT^ for the different experimental groups. For imaging experiments, significance was determined using a one-way ANOVA followed by Bonferroni’s multiple comparisons tests. For Ephys experiments, significance was determined using a two-tailed Student’s t-test. Where applicable, either p < 0.05 (∗), p < 0.001 (∗∗), or p < 0.0001 (∗∗∗) was considered a statistically significant difference.

### RF hyperpolarizes and inhibits the firing of neurons expressing TMEM16A^FeRIC^

In neurons, activation of a Cl^-^ current can either depolarize or hyperpolarize the membrane potential, depending on the difference between the Cl^-^ reversal potential and the resting membrane potential. The Cl^-^ reversal potential is set by the intra and extracellular Cl^-^ concentrations which are established by the interplay of diverse Cl^-^ channels and transporters (*37*). In mature neurons, the Cl^-^ reversal potential is usually negative with respect to the resting membrane potential. Then, the activation of a Cl^-^ channel produces Cl^-^ influx resulting in membrane hyperpolarization that leads to neuronal inhibition. To examine the RF-induced neuronal inhibition, we patch-clamped hippocampal neurons expressing TMEM16A^FeRIC^ to monitor the membrane potential and the spontaneous action potentials before RF (baseline), during RF (180 MHz and 1.6 μT), and within the 2-minute time window after RF stimulation (post-RF). We first verified that neurons expressing TMEM16A^FeRIC^ display similar electrical properties compared to mock-transfected neurons (**Fig. S1C-E**). Next, we tested the effects of RF stimulation on neurons expressing TMEM16A^FeRIC^. In those neurons, RF stimulation drove the membrane potential to more negative values with respect to the baseline (Baseline: -61.6 ± 2.2 mV; Post-RF: -64.1 ± 2.4 mV, n = 10 cells; p < 0.05) (**Fig. 6A, C**). This effect was inhibited with the TMEM16A antagonist Ani 9 (1 µM) (**Fig. 6B, D;** Baseline: -62.7 ± 1.7 mV, Post-RF: -62.5 ± 1.7 mV; n = 12 cells; p = 0.43). To evaluate the effect of RF on neuronal excitability, we recorded spontaneous action potentials before RF (baseline), during RF (180 MHz and 1.6 μT for 5 min), and in the post-RF period. RF consistently decreased the frequency of spontaneous action potentials in TMEM16A^FeRIC^-expressing neurons (Baseline: 0.0258 ± 0.0169 Hz, Post-RF: 0.0088 ± 0.0091 Hz; n = 8 cells; p < 0.05) (**Fig. 6E**), but RF did not affect those neurons treated with Ani 9 (Baseline: 0.0813 ± 0.0715 Hz, Post-RF: 0.0773 ± 0.065 Hz; n = 5 cells; p = 0.65) (**Fig. 6F**). Next, we evaluated if RF hinders the activation of neurons evoked by a strong depolarizing stimulus, such as high extracellular K^+^ concentration (70 mM). In neurons co-expressing gCaMP6 and TMEM16A^FeRIC^, high K^+^ produced robust Ca^2+^ transients (Max: 6.8 ± 0.5 ΔF/F0, AUC: 1503 ± 119, n = 21 cells) (**Fig. 7A, B**). The Ca^2+^ responses evoked by high K^+^ were similar to those evoked in mock-transfected neurons and neurons expressing either TMEM16A^FeRIC^ or TMEM16A^WT^ (**Fig. S4A, B**). However, treating neurons expressing TMEM16A^FeRIC^ with RF for 10 minutes before applying the high K^+^ stimulus produced significantly smaller Ca^2+^ transients (Max: 4.4 ± 0.5 ΔF/F0, AUC: 1061 ± 146, n = 38 cells, p < 0.05). This effect was prevented by Ani 9 (Max: 7.3 ± 1 ΔF/F0, AUC: 1500 ± 215, n = 9 cells, p < 0.05) (**Fig. 7A, B**). Conversely, in neurons expressing TMEM16A^WT^, stimulation with high K^+^ produced similar Ca^2+^ transients observed in non-stimulated neurons (Max: 9.1 ± 0.8 ΔF/F0, AUC: 1629 ± 196, n = 10 cells) and those stimulated with RF (Max: 9 ± 0.9 ΔF/F0, AUC: 2144 ± 263, n = 7 cells) (**Fig. S4C, D**). In a different series of experiments, we tested the inhibitory effects of RF in neurons expressing TMEM16A^FeRIC^ treated with bicuculline (5 µM), an antagonist of GABA_A_ receptors. Bicuculline increases the spiking activity by blocking the inhibitory GABAergic transmission (*38*). The neuronal spike activity was examined using BeRST 1. Neurons expressing TMEM16A^FeRIC^ treated with bicuculline for 10 minutes displayed robust spiking activity (0.15 ± 0.07 Hz, n = 44) (**Fig. 7C, E**) that was drastically decreased by the co-application of RF (**Fig. 7C– - E**) (0.002 ± 0.001 Hz, n = 29, p < 0.05). The RF effect on the spiking activity of TMEM16A^FeRIC^-expressing neurons was prevented by Ani 9 (0.28 ± 0.04, n = 20, p < 0.05) (**Fig. 7D**). Notably, the spiking activity produced by bicuculline in neurons expressing TMEM16A^FeRIC^ was significantly smaller compared with that produced in mock-transfected neurons (0.57 ± 0.14 Hz, n = 39, p < 0.0001), suggesting that TMEM16A^FeRIC^ expression itself decreases the neuronal excitability. Nonetheless, our results indicate that RF activates TMEM16A^FeRIC^, causing anionic influx in N2a cells and hyperpolarizing the membrane potential in neurons. Additionally, in neurons expressing TMEM16A^FeRIC^, RF stimulation hinders the activation of neurons evoked by high extracellular K^+^ or by blocking the GABAergic transmission.

**Fig. 6.**
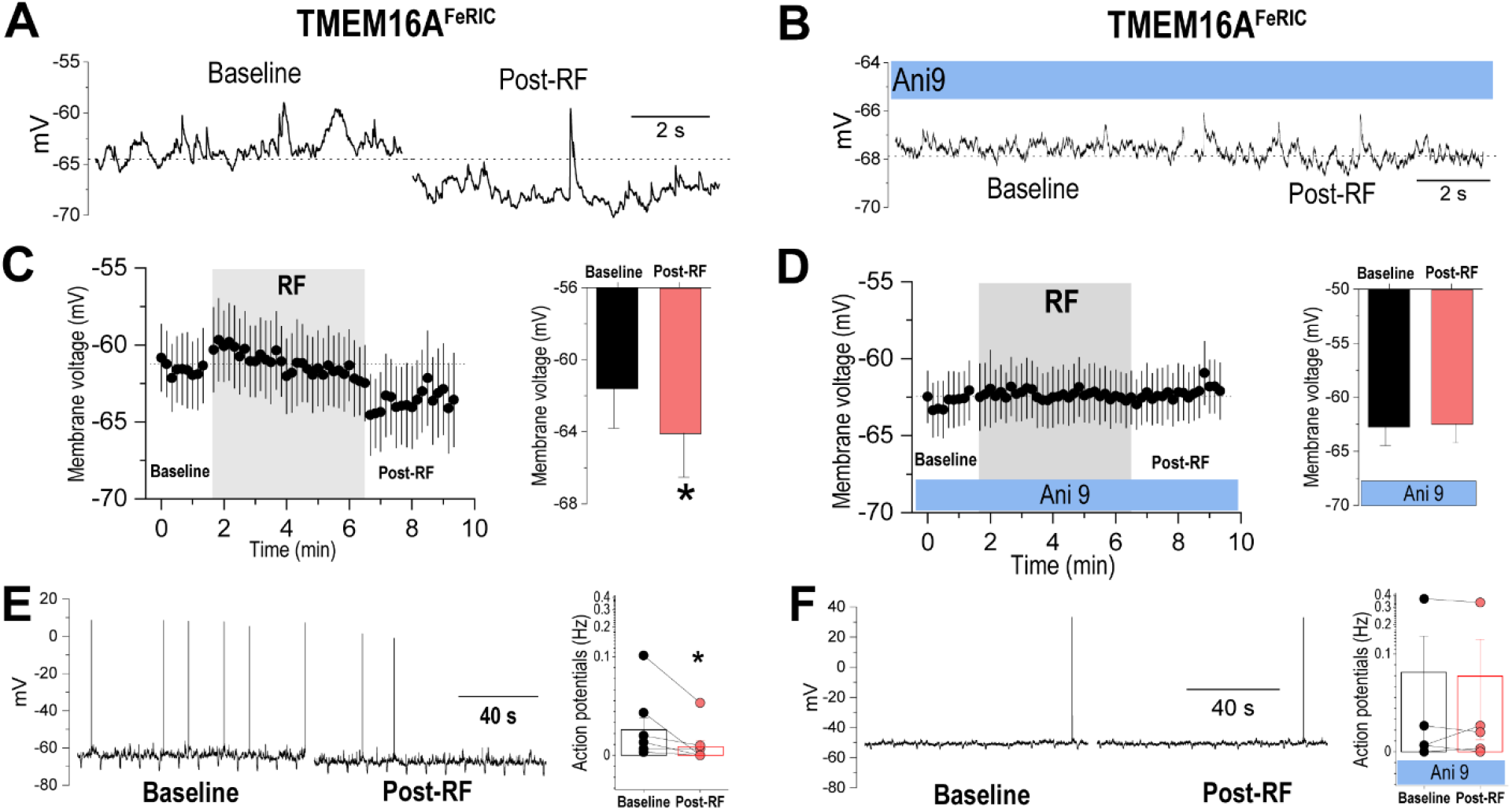
RF hyperpolarizes and inhibits neurons expressing TMEM16A^FeRIC^. (**A** - **D**) Membrane voltages recorded from neurons in the current-clamp mode. Representative traces of the membrane voltage in the baseline and within the 2 minutes after RF stimulation (post-RF) in neurons expressing TMEM16A^FeRIC^ (**A**) in the absence or (**B**) in the presence of the antagonist Ani 9. (**C, D**) Average membrane potential (±SEM) before RF, during RF, and in the post-RF period in neurons expressing TMEM16A^FeRIC^ (**C**) in the absence or (**D**) the presence of Ani 9. Insets: average membrane potential (±SEM) in the baseline and the post-RF period. (**E, F**) Spontaneous action potential firing in the baseline and the post-RF period in neurons expressing TMEM16A^FeRIC^ (**E**) in the absence or (**F**) in the presence of Ani 9. Insets: number of action potentials (±SEM) in neurons expressing TMEM16A^FeRIC^ in the baseline and the post-RF period. Significance was determined using a two-tailed Student’s t-test. Where applicable, p < 0.05 (∗) was considered a statistically significant difference.

**Fig. 7.**
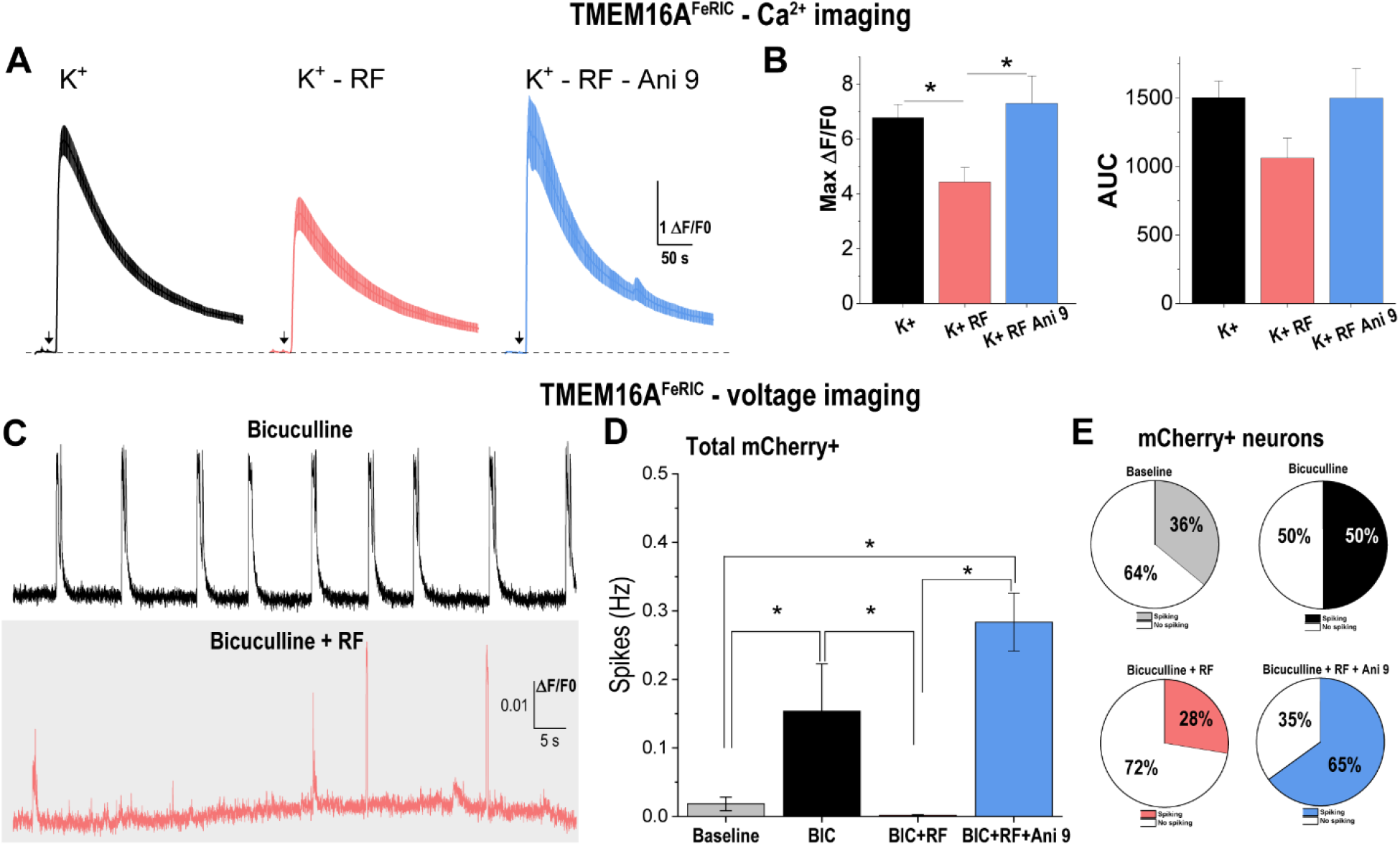
RF decreases the excitability of neurons expressing TMEM16A^FeRIC^. (**A**) Average changes (±SEM) in GCaMP6 fluorescence in neurons expressing TMEM16A^FeRIC^ following the addition of 70 mM K^+^ (black arrow) in non-stimulated (black), stimulated with RF (red) or stimulated with RF in the presence of Ani 9 (blue). (**B**) Bar graphs are the averages (±SEM) of the maximum change in GCaMP6 fluorescence (Max ΔF/F0) and the area under the curve (AUC) of neurons expressing TMEM16A^FeRIC^ for the different experimental groups. (**C**) Representative changes in BeRST 1 fluorescence in neurons expressing TMEM16A^FeRIC^ following treatment with bicuculline for 10 min in the absence (top) or the presence of RF stimulation (bottom). (**D**) Averages (±SEM) of the spiking frequency (Hz) for all mCherry+ neurons (spiking and non-spiking) for all the experimental groups. Experimental groups are independent samples. Baseline corresponds to the spike frequency in the absence of any stimulus. Neurons were imaged after treatment with bicuculline for 10 min in the absence or the presence of RF or RF plus Ani 9. (**K**) Pie charts of the fraction of mCherry+ neurons that were spiking (at least one spike in the entire experiment) or not spiking for the different experimental groups. Significance was determined using one-way ANOVA followed by Bonferroni’s multiple comparisons tests. Where applicable, either p < 0.05 (∗), p < 0.001 (∗∗), or p < 0.0001 (∗∗∗) was considered a statistically significant difference.

## Discussion

The magnetogenetic technique FeRIC allows the remote control of cell activity and neuronal excitability using RF magnetic fields. Here, we showed that FeRIC can control neurons by using the nonselective cation channel TRPV4 and the chloride permeable channel TMEM16A. In neurons expressing TRPV4^FeRIC^, RF depolarizes the membrane potential, decreases the membrane resistance, and increases the spiking frequency. Conversely, in neurons expressing TMEM16A^FeRIC^, RF hyperpolarizes the membrane potential and inhibits both the spontaneous and evoked spiking activity. Therefore, our results are consistent with the expected effects of activating TRPV4 and TMEM16A on the cells’ electrophysiological properties.

Several studies have reported the use of magnetic actuators, such as TRPV1 and TRPV4, coupled with ferritin to control cell activity using static and RF magnetic fields (*3–8, 10*). Because TRPV1 and TRPV4 are permeable to Ca^2+^, most studies have used Ca^2+^ imaging to monitor their activation. However, there was a lack of Ephys characterization of the activation of TRPV1 and TRPV4 using magnetic fields. This shortcoming, together with the controversy surrounding the mechanisms responsible for the magnetic activation of ferritin-coupled channels, has added uncertainty to the field of Magnetogenetics. Furthermore, whereas some magnetogenetic techniques, such as FeRIC, have reported an activation efficacy of about 20 – 40 % of cells, others, such as those using Magneto2.0, have not been reproducible when tested by independent groups (*19–21*). Here, we aimed to corroborate the fact that FeRIC is suitable for magnetically activating TRPV4^FeRIC^, producing the effects on the membrane’s electrical properties expected for this ion channel, which is permeable to Na^+^, K^+^, and Ca^2+^. Additionally, we examined the effects of activating the new TMEM16A^FeRIC^ channel with RF on the membrane potential. To examine the RF-induced activation of FeRIC channels, we conducted Ephys recordings combined with Ca^2+^ and voltage imaging, pharmacology, and genetic tools. Previously, corroborating the magnetic activation of ferritin-tagged channels with Ephys was precluded due to the incompatibility between the Ephys setup and the RF system. Unlike other Magnetogenetics methods that use near-static or kHz magnetic fields, FeRIC with a 180 MHz stimulus has inherent frequency isolation between the stimulus and the low-frequency kHz Ephys recordings, which reduces the electromagnetic interference. In our case, the stimulus is bandpass filtered by the 180 MHz resonant RF coil, and the Ephys circuitry contains low pass filtering as a standard component. Hence, the 180 MHz stimulus ideally should have no effect on the Ephys recordings. However, the Ephys setup and the RF system are not ideal in practice. While these filters greatly reduce the interference, nonlinearity in various components such as transistors or ferrous metals can still transform the 180 MHz signals into low frequency signals. In particular, second order nonlinearities can transform oscillating high-frequency signals into static signals, which explains the shift in the basal current levels observed when the RF is turned on. The 180 MHz signals induced on the Ephys circuitry turn into static shifts via non-linearities. Due to higher order effects, the conversions are very small, but not negligible when tiny pA currents and mV voltages are measured. To further reduce the electromagnetic interference between the Ephys setup and the RF system, a small RF emitting coil, attached at the base of the recording chamber, was used. While preserving the magnetic field strength at a patch-clamped cell of interest, a small (versus a large) coil is able to reduce the total electromagnetic interference on the much larger Ephys setup. However, since the maximum RF strength that did not produce noteworthy interference was about 1.6 µT, we were limited in testing stronger RF stimuli. To ensure that the coaxial cables connected to the RF coil did not act as antennas producing unwanted RF interference, two ferrite beads designed to block frequencies near 180 MHz were placed on the cables. From Maxwell’s equations, time-varying magnetic fields also produce time-varying electric fields. However, as discussed in our previous study (*5*), the values of these electric fields are too small for direct neuron stimulation. When an Ephys setup is used, voltages oscillating at 180 MHz are also induced directly on the patch-clamp circuit. These induced RF voltages are converted and scaled by the nonlinearities in the patch-clamp circuitry into baseline offsets, which should be minimized as much as possible. As shown in simulation and observed in practice, using RF at 180 MHz and a smaller RF coil combined with ferrite beads designed to block frequencies near to the RF stimulus frequency is a robust method to mitigate these offsets.

Solving the compatibility between the Ephys setup and the RF system allowed us to conduct Ephys recordings to corroborate the RF-induced activation of FeRIC channels, their biophysical properties, and the RF effects on the cells’ bioelectrical properties. The estimated unitary conductance for TRPV4^FeRIC^ was about 32 pS, which is in the range of the reported conductance values for wild-type TRPV4 (*29, 30*). Moreover, our results suggest that RF increases the TRPV4^FeRIC^ open channel probability, generating inward currents that are similar to those reported for native TRPV4 activated with endogenous ligands such as anandamide, arachidonic acid, and epoxyeicosatrienoic acid (*11*). The inward currents mediated by TRPV4 and TRPV4^FeRIC^ display a slow rising phase. Similarly, the deactivation phase is slow and lasts several minutes after the stimulus is removed (*4, 5, 11*). Remarkably, we previously reported that in cells expressing FeRIC channels, RF and ferritin interaction produces some of these endogenous actuators, which are responsible for TRPV4^FeRIC^ activation (*4*). It is important to note that the FeRIC technique is not suitable for instantaneously activating cells or neurons. RF gradually depolarizes the neuronal membrane potential and increases the spiking frequency of neurons expressing TRPV4^FeRIC^ on the minute scale. This effect may be explained by the fact that RF drives the neuronal membrane potential closer to the action potential firing threshold. Therefore, RF-induced membrane depolarization may have a significant influence on synaptic integration, neuronal excitability, and consequently on several neuronal functions, as we observed in increased spiking activity under RF stimulation. Our results contrast with previous studies that report immediate triggering of action potential firing in neurons expressing TRPV1 tagged with ferritin (*6, 7*) or TRPV4 fused with ferritin (*10*) upon stimulation with static magnetic fields. However, these reports did not provide further characterization of the magnetic activation of the ferritin-coupled TRPV1 and TRPV4 channels. Other groups reported that the magnetic stimulation of neurons expressing the ferritin-fused TRPV4, known as Magneto2.0, does not change any electrophysiological property in neurons. These differences might be explained by several factors such as how ferritin is tagged to the TRPV channels, the type of the magnetic stimulus, the expression level of the channels, among others.

In this study, we also reported the magnetic activation of the new TMEM16A^FeRIC^ channel, which allows hyperpolarization and inhibition of neurons. RF robustly decreases both the spontaneous and evoked neuronal spiking and Ca^2+^ activity. This observation is consistent with the role of homologous TMEM16B or Ano2, which controls the spontaneous and evoked action potential firing in cerebellar neurons (*39, 40*). Voltage imaging results provide evidence for the magnetic control of the FeRIC channels to manipulate the membrane potential in neurons and other cells. Therefore, FeRIC using TMEM16A^FeRIC^ offers the opportunity to inhibit neurons and control their action potential firing to interrogate their role in the neuronal circuitry.

A limitation of this study is that controlling neuronal excitability with the FeRIC technology is slow when compared to the millisecond-scale control offered by optogenetics. These slower dynamics can be attributed to the intrinsic TRPV4 activation kinetics seen when TRPV4 is activated by lipids, which attests to the complex biochemical mechanism that underlies Magnetogenetics. FeRIC is therefore suited for addressing problems that do not require manipulation of cells at a millisecond time scale, e.g., studying the effects of tonic transmission or plasticity in the functioning of the neuronal circuitry. The next steps that need to be taken to corroborate the utility of the FeRIC technique to control neuronal excitability is to conduct experiments *in vivo*.

The main advantages of FeRIC are the use of magnetic fields to activate ion channels and the utilization of endogenous ferritin as a RF transducer. Magnetic fields are noninvasive and penetrate virtually through the entire nervous system (*2*). In comparison, optogenetic approaches require invasive procedures for light delivery and are limited by light scattering and the light’s poor penetration into tissue (*41*). Furthermore, magnetic fields interact weakly with biological tissue, and thus avoid off-target effects, while optogenetics suffers from undesired increases in neuronal firing (*42*) that are caused by high-power light. High-power light raises the temperature in surrounding tissue (*43*), which, in turn, increases the neuronal firing (*42*).

In conclusion, we show that the FeRIC technique allows the control of neuronal bioelectrical properties. FeRIC technique opens opportunities to interrogate the relationship between neuronal excitability and diverse brain functions in freely moving experimental models.

## Materials and Methods

All animal experiments were approved by the Duke University Animal Care and Use Committee and by the UC Berkeley Animal Care and Use Committees and conformed to the NIH Guide for the Care and Use of Laboratory Animals and the Public Health Policy.

### Cell lines

Neuro2a cell line (N2a, ATC, CCL-13) was used. N2a cells were obtained from the UCB Cell Culture Facility (University of California Berkeley). Cell identity and negative Mycoplasma contamination were verified by the UCB Cell Culture Facility. Cells were maintained in Dulbeccos’s Modified Eagle Medium (DMEM, Gibco) supplemented with 10% fetal bovine serum (FBS, hyclone) and 100 units/mL penicillin, and 100 mg/mL streptomycin at 37°C and 5% CO_2_.

### Primary neuron cultures

Rat hippocampal neurons were obtained as previously described (*31, 32*). Briefly, hippocampi were dissected from embryonic day 18 Sprague−Dawley rats (Charles River Laboratory) and maintained in HBSS (Gibco #14170112). Hippocampal tissue was enzymatically treated with 2.5% trypsin (15 min at 37 °C). Next, the tissue was mechanically dissociated using a fire polished Pasteur pipette in minimum essential media (MEM, Invitrogen #11090-081) supplemented with 5% FBS, 2% B-27 (Gibco #17504044), 2% D-glucose, and 1% glutamax (Gibco # 35050061). Cells were plated onto Poly-D-lysine (1mg/mL, Sigma #P0899) coated coverslips (12 mm) at 6 × 10^4^ cells/cm^2^ in MEM supplemented medium. 24 hours after, 50% of the MEM supplemented media was replaced with Neurobasal medium (Invitrogen #21103-049) supplemented with 2% B-27 supplement and 1% glutamax. Cells were maintained at 37°C and 5% CO_2_. Experiments using cultured hippocampal neurons were conducted between 12 – 14 days after seeding because at this time, cultured neurons have established circuits (*44*).

### Plasmids

The constructs TRPV4^FeRIC^ and TRPV4^WT^ were obtained as described previously (*3*). To generate the TRPV4^WT^ construct, we used full-length rat TRPV4 cDNA, which was a gift from R. Lefkowitz (Duke University). Spe I and Not I restriction sites were introduced using PCR. The full-length wild-type TRPV4 was subcloned into the PLVX-IRES-mCherry vector to generate TRPV4^WT^ (Clontech, Catalog No. 631237). To generate the TRPV4^FeRIC^ construct, PCR primers were designed to eliminate the 3′ stop site in wild-type TRPV4 and introduce a 3′ Not I site. PCR primers introducing a 5′ Not I site, and a 3′ BamH I site and a stop codon were used to amplify human Kininogen1 domain 5 (FeRIC). This FeRIC fragment was subcloned into the Xba I and BamH 1 sites within the PLVX-IRES-mCherry vector containing TRPV4. To generate the TMEM16A WT construct, we used full-length TMEM16A cDNA, which was obtained from Origene (mouse untagged clone, PCMV6-Kan/Neo, #MC205263). TMEM16A cDNA was subcloned into the PLVX-IRES-mCherry vector between the 5’ NotI and 3’ BamH I sites. To generate TMEM16A^FeRIC^, PCR primers were designed to introduce the 5’ EcoRI, and the 3’ Not I and to eliminate the stop codon in the FeRIC (human Kininogen1 domain 5) fragment. This FeRIC fragment was subcloned into the EcoR I and Not I sites within the PLVX-IRES-mCherry vector containing TMEM16A. All completed constructs were sequence-verified by the Molecular Cell Biology Sequencing Facility (UC Berkeley) and analyzed using Serial Cloner 2.6 and Benchling.

### Chemical transfection of N2a cells

N2a cells were plated on non-coated glass-bottom 35-mm dishes or non-coated 12-mm coverslips placed into 24-well plates. Cells were seeded at a density of 3 × 10^4^ cells/cm^2^ and cultured in DMEM supplemented medium. After 18-24 hours, cells were transfected using the Lipofectamine LTX Plus reagent (ThermoFisher #15338030) with wild-type or FeRIC channels. For imaging experiments, N2a cells were co-transfected with the wild-type or FeRIC channels and GCaMP6 (GCaMP6 medium, Addgene cat.40754) or YFP-H148Q. For transfection, OptiMEM free serum medium (ThermoFisher #31985088) was used to prepare the DNA/Lipofectamine LTX mix. The transfection mix had the following composition per each 35-mm dish: 300 μL OptiMEM, 4 μL Lipofectamine LTX, 3 μL PLUS reagent, 0.7 μg ion channels DNA, and 0.7 μg of GCaMP6 DNA. The transfection mix had the following composition per 12-mm coverslip into 24-well plates: 100 μL OptiMEM, 1 μL Lipofectamine LTX, 1 μL PLUS reagent, 0.2 μg ion channels DNA, and 0.2 μg of indicators DNA.

### Chemical transfection of cultured neurons

Neurons plated on 12-mm coverslips were seeded at 7 × 10^4^ cells/cm^2^ and cultured in Neurobasal supplemented medium. After 8 - 14 days, cells were transfected using the Lipofectamine 2000 (ThermoFisher #11668019) with GCaMP6 (GCaMP6 medium, Addgene cat.40754) and the FeRIC channels. For transfection, OptiMEM free serum medium (ThermoFisher #31985088) was used to prepare the DNA/Lipofectamine 2000 mix. The transfection mix had the following composition per each coverslip/well: 100 μL OptiMEM, 2 μL Lipofectamine 2000, 0.5 - 0.7 μg FeRIC channels DNA, and 0.5 μg of GCaMP6 DNA. Neurons were incubated with the transfection mix at 37°C for three hours. Next, the culture medium containing the transfection mix was replaced with fresh Neurobasal supplemented medium. Experiments were conducted 3 – 4 days after transfection.

### RF coil

The RF emitting coil consisted of a single loop of wire with a loop diameter of about 2.8 or 5 cm. The coil was connected in parallel with tuning capacitors, forming an LC circuit resonant near 180 MHz. The circuit was matched to about 50 ohms near 180 MHz using a series capacitor. The coil was checked to still be matched near 180 MHz when parasitic capacitance due to nearby metal (e.g., from a microscope) was present. The RF signal was generated by a broadband (35 MHz to 4.4 GHz) signal generator (RF Pro Touch, Red Oak Canyon LLC) and amplified using a 5W amplifier (Amplifier Research, model 5U100, 500 kHz to 1000 MHz). The magnetic field produced by the coils was measured using EMC near-field probes (Beehive Electronics model 100B) connected to a spectrum analyzer (RSA3015E-TG, Rigol Technologies). The Beehive 100B probe was provided with magnetic field calibration data from the manufacturer and proper scaling factors for 180 MHz measurements were interpolated based on this data. The Rigol spectrum analyzer was manufacturer calibrated with a system registered to ISO9001:2008. Using the Beehive probe and the Rigol spectrum analyzer, the magnetic field strength was measured to be about 1.6 µT for 180 MHz at a location about 3 mm above the recording chamber. Unless stated otherwise, these are the values used throughout the manuscript. The measurements were made slightly above the cell culture dish location because the probes could not be lowered into the coils without changing the angle of the probes with respect to the magnetic field.

To ensure that the coaxial cables connected to the RF coil did not act as antennas producing unwanted RF interference, two ferrite beads designed to block frequencies near 180 MHz (Laird-Signal Integrity Products manufacturer part number 28A0592-0A2) were placed on the cables. This was especially important because without the ferrite beads, when the RF stimulus was turned on, the shift in basal current levels on the Ephys recordings increased by several orders of magnitude and large RF signals could be picked up in the air by a spectrum analyzer and a loop probe far away from the RF coil.

### Measurements of the voltage or current basal shift produced by an RF source

RF voltages induced on the headstage are transmitted into the Ephys recordings in the form of a baseline shift. The baseline shift was measured as a function of the RF magnetic field produced by a 2.8 cm diameter coil pipette under in-the-bath conditions or as a function of RF voltage produced by a Siglent SDG6022X arbitrary waveform generator directly connected (for the current-clamp configuration) or capacitively coupled (for the voltage-clamp configuration) to the headstage. In the case of direct/capacitively coupled connections, a voltage source and a series 50 Ω resistance, corresponding to the waveform generator and its output impedance, respectively, were connected directly to the headstage. A 50 Ω characteristic impedance cable of about 2 feet was between the waveform generator and the headstage. Given the 50 Ω output impedance, the power and thus voltage delivered to the headstage is not dependent on the length of the cable, except for minor losses negligible for short cable lengths. For voltage-clamp measurements, in order to present a large low-frequency impedance to the headstage and prevent the headstage from attempting to provide large currents, a 10 pF capacitor (Johanson Technology 102S42E100JU3S) was placed in series between the headstage and the cable immediately next to the headstage. This capacitance passes the RF from the waveform generator to the headstage while blocking static currents between the waveform generator and the headstage.

Using a Multiclamp 700A amplifier (Axon Instruments), we measured the baseline shift in pA for the voltage-clamp configuration or the baseline shift in mV for the current-clamp configuration as a function of RF voltage on the voltage source from the waveform generator. The data were fit with a quadratic term or higher order terms for larger voltage values.

### Ca^2+^ imaging

Epifluorescence imaging experiments were conducted as previously described (*4, 5*). Cytosolic levels of Ca^2+^ were monitored by fluorescence imaging of cells positive for GCaMP6. Cells expressing FeRIC channels were identified as those cells with mCherry^+^ expression. Experiments were conducted using an upright AxioExaminer Z-1 (Zeiss) equipped with a camera (Axiocam 702 mono) controlled by Zen 2.6 software. Excitation light was delivered from a LED unit (33 W/cm2; Colibri 7 Type RGB-UV, Zeiss). mCherry was excited at 590/27 nm and emission was collected at 620/60 nm. GcaMP6 was excited at 469/38 nm and emission was collected at 525/50 nm. Illumination parameters were adjusted to prevent overexposure and minimize GcaMP6 photobleaching. All experiments corresponding to a series were done under the same illumination settings. Images were captured with a W “Plan-Apochromat” 20x/1.0 DIC D=0.17 M27 75mm lens at 1 frame/s. Experiments were carried out at room temperature (20 – 22 °C) using HBSS (Invitrogen, 14025092). At the beginning of imaging experiments, cells were washed three times with 1 mL of the HBSS. Next, the dish was placed onto the microscope stage and the cells were localized with transmitted illumination (bright-field). Next, with reflected illumination, the fluorescence signals from mCherry and GcaMP6 were corroborated, and the field of interest was selected. Preferred fields were those with isolated and healthy cells. A thermocouple coupled to a thermistor readout (TC-344C, Warner Instruments) was placed inside the plate in direct contact with the HBSS. The temperature of the HBSS was monitored during the experiment (Temperature initial: 22.09°C; Temperature final: 22.03°C; ΔT: -0.6°C; n=305). Cells were rested for about 10 minutes before imaging. RF was delivered using a custom-built RF-emitting coil. Cells were stimulated with RF fields at 180 MHz (at 1.6 µT). For each experiment, cells were imaged for the first 60-120 s with no stimulus (Basal) and followed with RF exposure for 4-6 min. After RF stimulation, cells were exposed to GSK101, which is a TRPV agonist, by pipetting 1 mL of the GSK101 diluted in HBSS (at 2μM) to reach the final 1μM concentration.

### Halide imaging – YFP-H148Q (YFP)

Cytosolic levels of iodide were monitored using fluorescence imaging of cells positive for YFP. Cells expressing TMEM16A^FeRIC^ or TMEM16A^WT^ were identified as those cells with mCherry^+^ expression. Experiments were conducted using the same microscope system as that used for Ca^2+^ imaging. The experiments were carried out at room temperature (20 – 22 °C) and cells were maintained in Live Cell Imaging Solution (Invitrogen, A14291DJ) that contains 140 NaCl,

2.5 KCl, 1 MgCl_2_, 1.8 CaCl_2_, 20 HEPES, 5 mM glucose, pH 7.4. Cells were stimulated for 9 minutes with either RF or ionomycin before the time-lapse image acquisition. Next, fluorescence images of YFP were acquired (1 frame/s) for 1 minute (basal line). Then, 1 mL of iodide-containing imaging solution was added to reach a final concentration of 70 mM iodide. Fluorescence of YFP was continuously imaged for 5 minutes. Iodide solution contained (in mM): 140 NaI, 5 NaCl, 2.5 KCl, 1 MgCl_2_, 2 CaCl_2,_ 5 mM Glucose, 20 HEPES, pH 7.3. In the second series of experiments, cells were stimulated for 10 min with the RF or ionomycin, but simultaneously with Ani 9 (1 µM).

### Voltage imaging – BeRST 1

Neurons previously transfected with either FeRIC or wild-type channels were incubated with an Imaging Solution (Gibco) containing BeRST (500 nM) at 37 °C for 20 minutes. Neurons expressing the FeRIC or wild-type channels were identified as those positive for mCherry. Fluorescence imaging of BeRST 1 was conducted on the AxioExaminer Z-1 microscope described above. Images were acquired with a W-Plan-Apo 20×/1.0 water objective (20×; Zeiss) at 1 frame/20ms. mCherry images were acquired as described above. BeRST 1 was excited at 640/30 nm and emission was collected at 690/50 nm. Images were analyzed using ImageJ (National Institutes of Health). Regions of interest (ROIs) were placed over the soma of neurons. Responses are presented as ΔF/F0, where F0 is the resting fluorescence averaged over 100 first frames at the start of acquisition. Customized code (MATLAB) was developed to analyze data and automatically count the fast changes of fluorescence of BeRST 1.

### Electrophysiological (Ephys) recordings

The electrical properties of cells were examined using the patch-clamp technique in the whole-cell configuration. Cells were transferred to a recording chamber filled with extracellular saline solution and placed on the microscope stage (AxioExaminer Z-1, Zeiss). N2a cells were recorded using Hanks’ Balanced Salt Solution (HBSS) (Thermo Scientific™ 14025-092) and cultured neurons were recorded using Live Cell Imaging Solution (Invitrogen™ A14291DJ) supplemented with 5 mM glucose. Whole-cell recordings were performed from the soma of N2a or hippocampal neurons with patch pipettes (4–8 MΩ), filled with an internal solution containing in mM: 135 KmeSO_4_, 10 KCl, 10 HEPES-K^+^, 5 NaCl, 5 ATP-Mg^2+^, 0.4 GTP-Na^+^, and pH 7.2–7.3.

For those extra and intracellular solutions, the chloride equilibrium potential was −58 mV. For experiments conducted to examine neurons expressing TMEM16A, the patch pipettes were filled with a low chloride intracellular solution containing in mM: 145 KmeSO_4_, 10 HEPES-K^+^, 3 NaCl, 5 ATP-Mg^2+^, 0.4 GTP-Na^+^, pH 7.2–7.3. The chloride equilibrium potential for the low chloride intracellular solution was −98 mV.

The voltage-clamp recordings were performed using a Patch-Clamp PC-501A (Warner Instruments Corp) or a Multiclamp 700A amplifier (Axon Instruments). In voltage-clamp experiments, the voltage holding (Vh) was adjusted to -60 mV and the capacitive currents were compensated up to 70%. For N2a cells, an experiment was accepted if the seal resistance was >1 GΩ and the access resistance (Ra; ∼10-30 MΩ) did not change > 20% during the span of the experiment. The Ra was monitored by applying a voltage step (−5 mV, 200 ms). The current elicited by the -5 mV voltage pulse was measured to calculate the membrane resistance (Rm). To obtain the voltage- vs-current (V-I) relation in N2a cells recorded in voltage-clamp mode, 12 voltage steps (1-s duration, -80 to -25 mV, ΔV = 10 mV) were applied. For neurons recorded in current-clamp mode, the membrane potential was set by injecting current and the Rm was monitored during the experiments with a current pulse (−5 pA, 500 ms). To obtain the current-vs-voltage (I-V) relation in neurons, 12 current pulses (1-s duration, -20 to 80 pA, ΔV = 10) were applied. To assess the effects of RF on neuron electrophysiological properties, I-V relations were characterized in resting conditions and after 5 minutes of RF stimulation. Data were acquired with a low pass filter at 3.0 kHz and sampled at 10.0 kHz using an A/D converter Digidata 1320A (Axon Instruments) or Axon Digidata 1550B (Molecular Devices). Digidata 1320A with pClamp 10.7 software (Molecular Devices) and Axon Digidata 1550B with pClamp 11.2 software (Molecular Devices) were used to acquire data and generate the current pulses and voltage steps.

### Quantification and statistical analysis

#### Simulations of the magnetic and electric fields produced by RF

The distributions of the electric (E) and magnetic (B) fields applied to the cells were simulated using the finite-difference time-domain (FDTD) method implemented by the openEMS project (https://openems.de/start/). Simulations were done considering the 5 cm-diameter RF coil or the 2.8 cm-diameter RF coil containing a recording chamber filled with imaging saline solution. The saline solution was geometrically modeled to reflect the two-compartment chamber used for Ephys recordings, and the relative permittivity and conductivity of the saline solution were set to 80 and 1.5 S/m respectively. Wires for the Ephys setup were modeled as perfect electrical conductors and the head stage input was modeled as a 500 MΩ resistance. Simulated voltage pulses were injected across the coils and the resulting time-domain fields and voltages were transformed into frequency-domain fields and voltages. At 180 MHz, all fields and voltages were scaled to give a 1.6 µT magnetic field amplitude at the bottom of the center of the recording chamber center. Given this magnetic field strength, amplitudes for electric fields at 180 MHz at the bottom of the recording chamber center and voltages at 180 MHz across the 500 MΩ head stage resistance were recorded. Cells and more complicated patch-clamp circuitry were not modeled.

#### Ca^2+^ imaging analysis

Cytosolic Ca^2+^ levels were monitored in cells expressing GCaMP6 and either FeRIC or wild-type TRPV channels. The fluorescence intensity of GCaMP6 was acquired at 1 Hz. GCaMP6 fluorescence was computed in a cell-based analysis with a customized MATLAB (Release 2018b, MathWorks Inc., Natick, Massachusetts) code. Maximum intensity projection was performed along the time axis to get the maximum intensity signal of cells expressing GCaMP6 in the field of view. In experiments where cells were stimulated with agonists, the maximum intensity projection corresponds to those cells that expressed functional ion channels. The watershed algorithm (MATLAB implemented function: Watershed transform) (*45*) was used to identify and label the cells to generate a cell-based mask for each experiment. The algorithm does not contain a motion correction component because the spatial movement of the cells during the time-lapse acquisition was negligible. The GCaMP6 fluorescence intensity was measured for each masked cell of the time-lapse acquisition (600 s). GCaMP6 fluorescence signal is presented as ΔF/F0, where F0 is the basal fluorescence averaged over 1-121 s before the start of stimulation and ΔF is the change in fluorescence with respect to the basal values. For analysis, GCaMP6 fluorescence measurements corresponding to the first 5 frames were discarded because of an inconsistent artifact. For each masked-cell, the data from 6-121 s were fit with a mono-exponential curve [*f*(*t*; *a, b*) = *ae*^*bt*^]. The fitted curve was used to correct the GCaMP6 photobleaching effect over the entire acquisition period. The masked cells that showed abnormal behavior, observed as the fitted growth factor (b) value above 0.002, were excluded from the analysis. Masked cells were considered responsive when the averaged ΔF/F0 over the time during the period from 122 to 601 s (RF-stimulation) increased 10 times over the standard deviation of the GCaMP6 ΔF/F0 of the basal session. For each experimental group, the change in GCaMP6 ΔF/F0 and the corresponding AUC (t = 121 – 601 s) of all analyzed masked-cells, which corresponded to all TRPV-expressing cells, were averaged. The plots of the GCaMP6 ΔF/F0 changes correspond to the data obtained from all identified cells, including both responsive and non-responsive cells. The MATLAB code that we used for the Ca^2+^ imaging analysis is available in Supplementary Materials.

#### Nonstationary noise analysis - Estimation of unitary current and conductance

The nonstationary noise analysis of macroscopic currents allows us to estimate diverse ion channels’ properties such as the unitary current (*i*). This analysis uses the relationship between mean current amplitude (*I*) and the variance of the current (σ^2^) given by 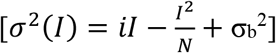. In this expression, N is the estimated number of channels and σ ^2^ is the background noise (*24, 25, 27, 28, 46*). A parabola is fitted for experimental data where the open-channel probability (P_o_) ranges from 0 to 1, but for P_o_ values smaller than 0.2, only the parabola roots are observed. The slope of in the neighborhood of either root of the parabola corresponds to *i*. For each inward current, the mean *I* amplitude and σ^2^ were measured. Next, the *I* σ^2^ values were used to obtain the *I*-σ^2^ relationship. In our conditions, we observed only the parabola root (Fig. 1 E), which was used to estimate *i*. The *i* value was used to estimate the conductance (γ) of TRPV4^FeRIC^ using the relationship given by 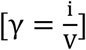, where V is the driving force that corresponds to the difference between the holding voltage (−60 mV) and the theoretical reversal potential for TRPV4 (5 mV).

#### Voltage imaging analysis – BeRST 1. Fast depolarization events/spiking rate inference

The slow imaging rate of the camera system used to observe the BeRST 1 fluorescence changes renders resolving single action potential firing challenging and difficult. The imaging system integrates BeRST 1 fluorescence signals over the sampling interval of 21.5 milliseconds. Because the observed BeRST 1 fluorescence signal is the result of fired action potentials convolved with an unknown BeRST 1 fluorescence signal integration kernel, the state-of-the-art deconvolution algorithm can be employed to infer the precise location of the action potentials underlying the integrated BeRST 1 time series data. We exploit an extension of the fast nonnegative deconvolution (FOOPSI) method (*47*). The constrained deconvolution approach permits the estimation of the general autoregressive time constants of varying orders, which allows the calculation of the transfer function that characterizes the transformation from neuronal spikes to the integrated BeRST 1 fluorescence changes data (*48*). Implementation of this constrained FOOPSI algorithm is found inside the deconvolution package of CaImAn, an open source calcium imaging data analysis tool (*49*). The constrained FOOPSI algorithm solves a convex optimization inverse problem with a sparsity prior assumption in the forward convolution model. The forward model of the algorithm assumes that the occurrences of spikes are sparse in time, which can be safely assumed. BeRST 1 ΔF/F0 data with the low frequency drift component removed was inputted into the algorithm with a custom noise standard deviation parameter and a custom baseline fluorescence value computed for each cell in every experiment. All other constrained FOOPSI algorithm parameters, including the unknown time constants of the kernel, are estimated automatically. The algorithm used CVX (http://cvxr.com/cvx/), a package used to solve convex programs (*50*).

The noise standard deviation parameter was computed for each BeRST 1 ΔF/F0 recording. For every neuron, the time series component corresponding to basal activity was computed using a moving standard deviation of length 23 samples, which equates to about 500 ms in time. Basal activity time series was taken to be the 23 samples-long vector that had the standard deviation equal to the median of the bottom 25^th^ percentile range of moving standard deviations. This process allowed us to automatically select the temporal range in the data corresponding to no observed spiking activity. Remaining components of the time series were taken to be the nonbasal activity time series. The noise standard deviation and the baseline fluorescence value parameters were set to the standard deviation and mean of this estimated basal activity component for each cell, respectively.

The constrained FOOPSI algorithm outputs inferred spikes of varying strengths given the above BeRST 1 ΔF/F0 data input and the noise standard deviation and baseline fluorescence value input parameters. Because inferred spikes of small amplitudes are often extraneous and artifacts of deconvolution, we developed a custom filtering routine to separate the most probable spikes from noise. In recordings where noticeable spikes were present, i.e., time series data where the percent change from the basal activity standard deviation to the nonbasal activity standard deviation exceeded 75%, only inferred spikes with amplitudes greater than the custom threshold of the baseline fluorescence value + 1.5*noise standard deviation were registered as components of the final inferred depolarization events. In order to mitigate extraneous, noisy spike events from biasing our inference of depolarization events, we increased this threshold to be the baseline fluorescence value + 4*noise standard deviation in recordings where there did not exist noticeable spikes in the BeRST 1 fluorescence data. This filtering algorithm pipeline, which is a slight modification of the thresholder FOOPSI algorithm, removes multiple false positive spikes, and yielded accurate and reliable inference of fast membrane depolarization events or spikes hidden under the integrated BeRST 1 fluorescence change data (*51*).

#### Statistical analysis

For all experimental groups, at least three independent experiments were conducted. Differences in continuous data sets were analyzed using Microcal OriginPro 2021 9.8 software (OriginLab). Data are means ± SEM. When only two experimental groups were compared, the statistical probe applied was the Student’s t-test. For comparisons of repeated-measurements, (basal, RF, post-RF), the two-tailed Student-test probe was applied. For results with normal distributions, hypothesis testing was performed using a one-way ANOVA, followed by the Bonferroni’s multiple comparisons test. For results that did not follow a normal distribution, the hypothesis testing was performed using a non-parametric one-way Kruskal-Wallis ANOVA, followed by Dunn’s multiple comparisons test. To compare results according to the levels of two categorical variables we used a two-way ANOVA, followed by Holm-Sidak multiple comparisons test. Where applicable, p < 0.05 (∗), p < 0.001 (∗∗), or p < 0.0001 (∗∗∗) was considered statistically significant differences.

## Supporting information

FeRIC_SI

## Acknowledgments

We thank Jingjia Chen (University of California, Berkeley) for developing the customized MATLAB code for quantifying the changes in YFP fluorescence.

## Author contributions

C.L. and E.J.B. conceived the FeRIC technology. C.L., M.H.-M., and K.W-M. designed the project. M.H.-M. designed and conducted the imaging experiments and analyses. V.H. built the RF coils. K.W-M. and V.H. implemented the Ephys and RF set up. K.W-M. designed and conducted the Ephys experiments and analyses. S.M.H. coded and analyzed the BeRST 1 data. K.P. and E.J.B. designed and cloned TMEM16A^FeRIC^. R.H.K. contributed to implementing Ephys recordings. E.M. provided the cultured hippocampal neurons and BeRST 1. M.H.-M., K.W-M., V.H, and C.L. curated the results and data analysis. M.H.-M. wrote the manuscript. C.L., K.W-M., S.M.H, V.H., R.H.K., E.M., and E.J.B. revised and edited the manuscript. All authors approved the manuscript.

## Data and materials availability

All data are available in the main text or the supplementary materials. The codes generated during this study are available at GitHub: https://github.com/LiuCLab/FeRIC. The cDNA sequence of TRPV4^FeRIC^ is available at GenBank with the identifier TRPV4^FeRIC^: MT025942 (https://www.ncbi.nlm.nih.gov/nuccore/). The statistical analysis, the Ephy and Ca^2+^ imaging analysis files are available at Mendeley Data (doi:10.17632/y3gbz9s9rp.1).

