## Supplementary material for "Controlling the bioelectrical properties of neurons with ferritin-based Magnetogenetics": FeRIC_SI

Miriam Hernández-Morales *et al.*

**This PDF file includes:**

Fig. S1. Expression of TRPV4$^{FeRIC}$ or TMEM16A$^{FeRIC}$ does not significantly affect the bioelectrical properties of N2a cells or hippocampal neurons.

Fig. S2. RF evokes inward currents and $Ca^{2+}$ transients in Neuro2a cells expressing TRPV4$^{FeRIC}$.

Fig. S3. Diversity of spike firing and bursting in hippocampal neurons expressing TRPV4$^{FeRIC}$ upon RF stimulation.

Fig. S4. Expression of TMEM16A$^{FeRIC}$ or TMEM16A$^{WT}$ does not affect the high $K^+$-induced $Ca^{2+}$ transients in hippocampal neurons.

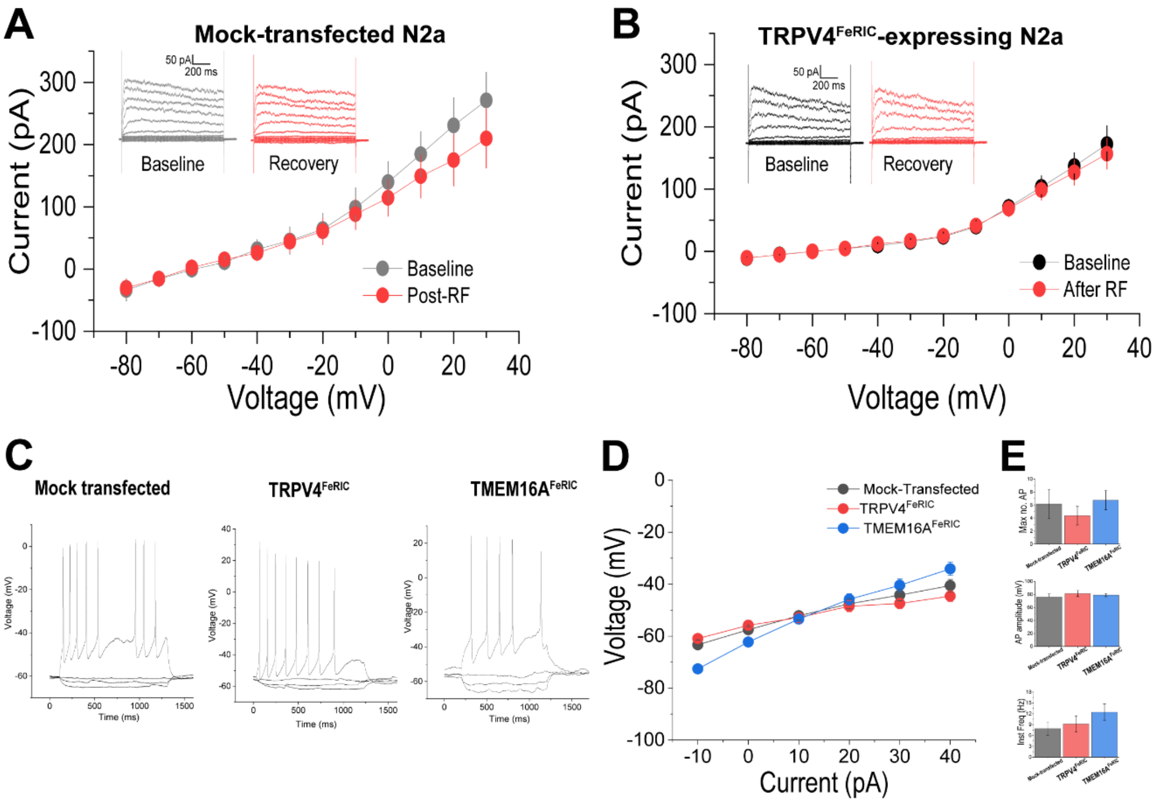


**Fig. S1. Expression of TRPV4$^{FeRIC}$ or TMEM16A$^{FeRIC}$ does not significantly affect the bioelectrical properties of N2a cells or hippocampal neurons. Related to Fig. 2 - 6.** (**A**, **B**) Voltage-currents (V-I) relations were obtained from (**A**) mock-transfected N2a and (**B**) N2a cells expressing TRPV4$^{FeRIC}$. Membrane currents were elicited by applying square voltage steps (starting at -80 mV with 10 mV increases) in cells recorded in the voltage-clamp mode. Insets: representative membrane currents generated by the square voltage pulses in the baseline (gray circles) and the 5 minutes after RF stimulation (recovery, red circles). (**C**) Voltages and action potentials (AP) generated by square current pulses (starting at -10 pA with 10 pA increases) applied to neurons recorded in the whole-cell current-clamp mode. Mock-transfected neurons (left), neurons expressing TRPV4$^{FeRIC}$ (middle), and neurons expressing TMEM16A$^{FeRIC}$ (right). (**D**) Current-voltage relations in mock-transfected neurons, neurons expressing TRPV4$^{FeRIC}$, and neurons expressing TMEM16A$^{FeRIC}$. (**E**) Average (±SEM) of active neuronal properties of mock-transfected neurons and neurons expressing TRPV4$^{FeRIC}$ or TMEM16A$^{FeRIC}$. Significance was determined using a one-way ANOVA followed by Holm-Sidak's multiple comparisons test.

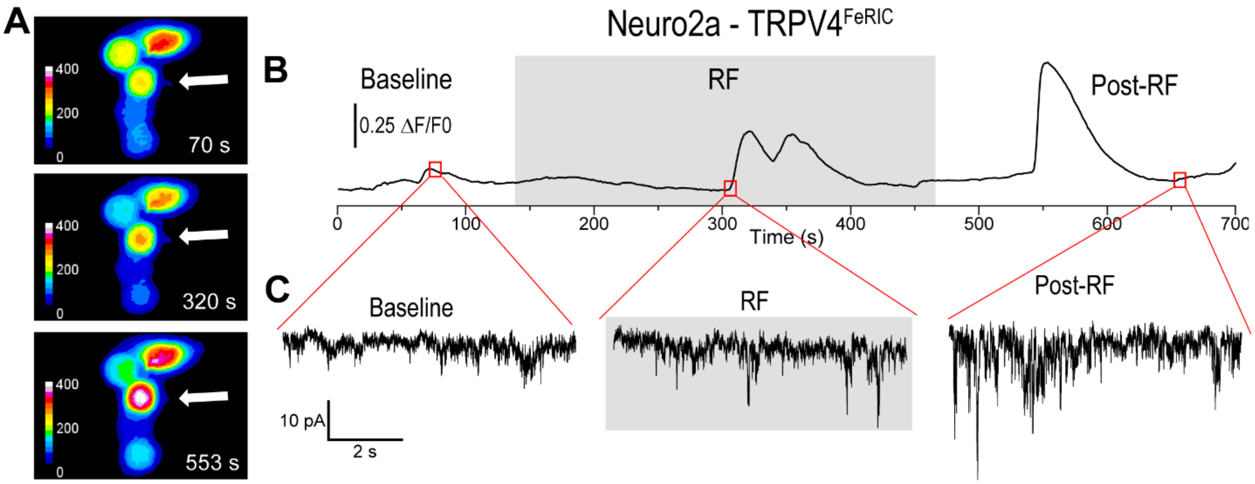


**Fig. S2. RF evokes inward currents and $Ca^{2+}$ transients in Neuro2a cells expressing TRPV4$^{FeRIC}$. Related to Fig. 2.** (**A**) Pseudocolor images of GCaMP6 fluorescence from N2a cells expressing TRPV4$^{FeRIC}$ before (70 s), during (320 s), and after RF stimulation (553 s). The white arrow indicates a patch-clamped N2a cell that was responsive to RF stimulation. (**B**) Changes in GCaMP6 fluorescence and (**C**) membrane currents from the cell in (A) before (baseline), during (gray box), and after RF (post-RF) stimulation.

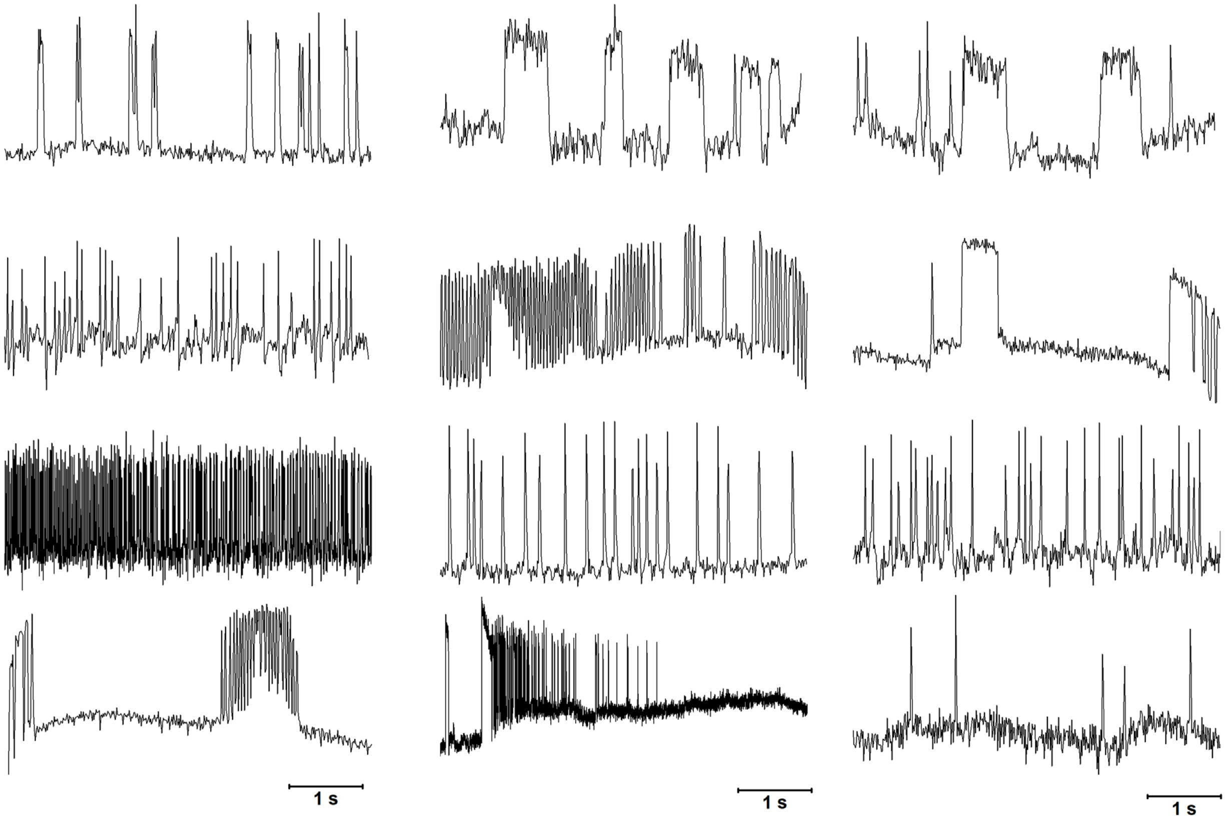


**Fig. S3. Diversity of spike firing and bursting in hippocampal neurons expressing TRPV4$^{FeRIC}$ upon RF stimulation. Related to Fig. 4.** Changes in BeRST 1 fluorescence in mCherry$^{+}$ neurons expressing TRPV4$^{FeRIC}$ following exposure to RF for 3 min.

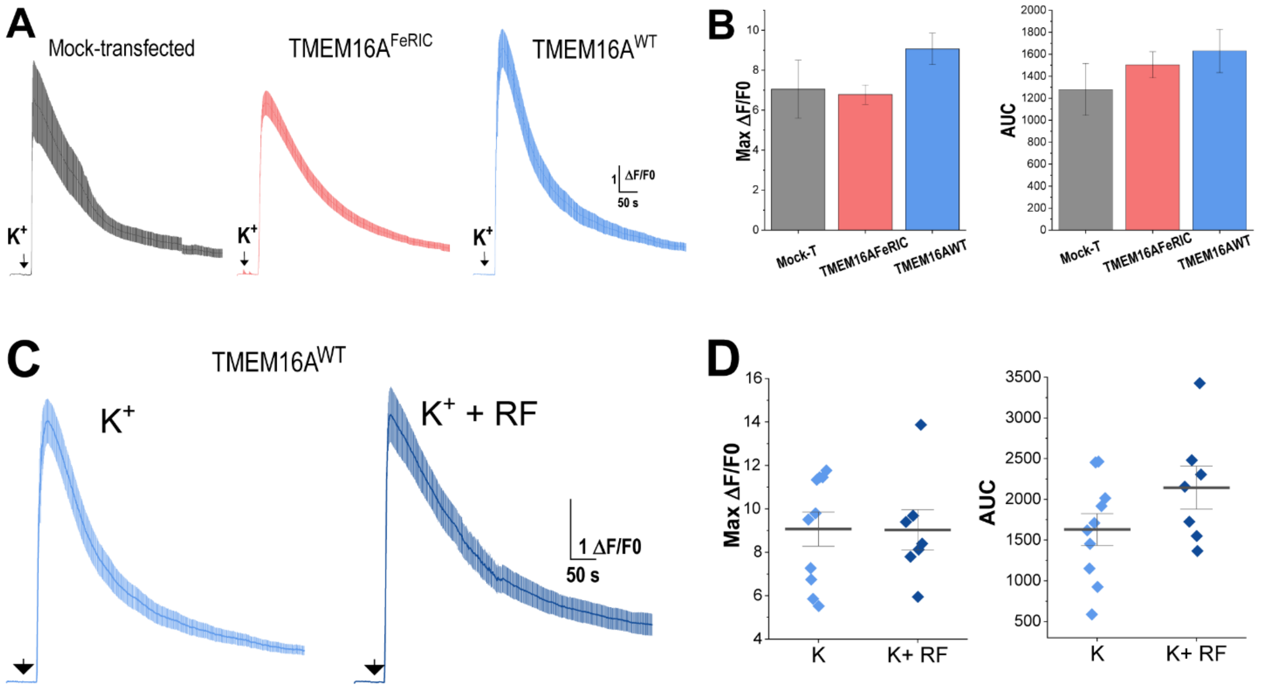


**Fig. S4. Expression of TMEM16A$^{FeRIC}$ or TMEM16A$^{WT}$ does not affect the high K$^{+}$-induced Ca$^{2+}$ transients in hippocampal neurons. Related to Fig. 7.** (**A**) Average changes in GCaMP6 fluorescence (±SEM), (**B**) the maximum change in GCaMP6 fluorescence (Max ΔF/F0), and the area under the curve (AUC) observed in mock-transfected neurons or neurons expressing TMEM16A$^{FeRIC}$ or TMEM16A$^{WT}$ following the addition of 70 mM K$^{+}$. (**C**) Average changes in GCaMP6 fluorescence (±SEM) in neurons expressing TMEM16A$^{WT}$ following the addition of 70 mM K$^{+}$ in the absence or upon RF stimulation. (**D**) Max ΔF/F0 and AUC in neurons expressing TMEM16A$^{WT}$ following the addition of 70 mM K$^{+}$ in the absence or upon RF stimulation. Significance was determined using a Student's t-test (2 experimental groups) or a one-way ANOVA followed by Bonferroni's multiple comparisons test (3 experimental groups).
